# Changes in gene regulation are associated with the evolution of resistance to a novel parasite

**DOI:** 10.1101/2025.08.13.670117

**Authors:** Lauren E. Fuess, Amanda K. Hund, Mariah L. Kenney, Meghan F. Maciejewski, Joseph M. Marini, Daniel I. Bolnick

**Affiliations:** Department of Biology, Texas State University, San Marcos, TX, USA; Department of Ecology and Evolutionary Biology, University of Connecticut, Storrs, CT, USA; Department of Biology, Carleton College, Northfield, MN, USA; School of Integrative Biology, University of Illinois at Urbana-Champaign, Urbana, IL, USA

## Abstract

Host-parasite interactions are ubiquitous and are important drivers of diversification and evolution. Host immune systems in particular are frequent targets of parasite-driven selection. The resulting rapid evolution of immune genes is usually framed as an ongoing ‘arms race’ between a co-evolving pair of host and parasite species. But, often immune evolution may be driven by the acquisition of a new and unfamiliar parasite. For instance, when marine populations of threespine stickleback (*Gasterosteus aculeatus*) colonized freshwater lakes ∼12,000 years ago, they encountered and evolved resistance to a freshwater-restricted cestode *Schistocephalus solidus*. We compared the transcriptomic responses of lab-reared stickleback from three populations with varying cestode susceptibility (naïve marine, susceptible lake, and resistant lake), when exposed to several immune stimuli. The resulting changes in expression reveal strong evidence for shared and population-specific responses during evolution of defense against a new parasite. Our investigation highlights the roles of several key immunological processes in underlying a general physiological response to tissue damage (fibrosis), and the importance of regulation of this fibrosis as a necessary step for its co-option into parasite defense. Furthermore, we highlight changes in expression of fibrosis-associated genes which facilitate faster and more targeted deployment of this defense against parasites; fish from resistant populations have higher and more consistent expression of fibrosis genes, which experience strong negative feedback in response to cestode stimuli only. Combined, our results provide strong evidence that changes in gene regulation and increased negative feedback to mitigate immunopathology are essential steps in the evolution of novel parasite defense.

## INTRODUCTION

The vertebrate immune system includes many evolutionarily ancient genes and biological processes (1). For instance, the NF-kB signaling pathway, central to innate immunity, likely originated in the common ancestor of metazoans or even earlier, and many of its functions are highly conserved across animals (2). Juxtaposed against this evolutionary conservatism, we also know that parasites can impose strong natural selection on host immune systems even over short time scales (3, 4). Selective forces induced by host-parasite interactions were first described in Van Valen’s Red Queen’s Race theory (5), and are widely studied (6–10). Parasitism has been shown to drive substantial immune diversification in animals, and some immune genes are among the fastest-evolving genes in animals (11, 12). How are we to resolve this apparent contradiction between evolutionary conservatism, versus rapid evolution, of animals’ immune systems? One possible explanation is that contemporary microevolution might rely on minor modification or co-option of existing genes and pathways (13–15), an example of the ‘evolutionary tinkering’ proposed by Jacob in 1977 (16). Therefore, an important question in evolutionary immunology is: Does animal adaptation to parasites mostly involve gain of new immune pathways, loss of old pathways, or co-option of existing pathways for new purposes? This question is particularly relevant to situations where a host population encounters a novel parasite species (as opposed to the long-term coevolution that is the focus of the Red Queen theory). Do hosts evolve immunity to a novel parasite through novel immune genes or gene regulatory pathways; or through co-option of existing genes or regulatory pathways?

The evolution of new immune defenses is not simply a gain of function, however. Animals must also suppress a defense when not required, and de-escalate after a successful response. Parasite defenses can be costly in a multitude of ways; excessive immune responses can cause host damage (i.e. “immunopathology”, (17–19)), and immunity is energetically costly, drawing resources from other beneficial processes (e.g., reproduction and growth rate; (20–22)). Natural selection does not typically favor ever-greater resistance, but rather an optimal balance between immune costs and benefits (23). Because of this balancing selection, adaptation to new parasites could drive either the evolution of up- or down-regulation of immune genes in a tightly coordinated series of events, and might promote either gain or loss of genes or pathways. In this paper, we report the results of a study to evaluate the extent to which adaptation to a new parasite involves gain versus loss, up-versus down-regulation, and co-option or novelty of immune gene expression and pathways in a model host-parasite system.

Recent studies have identified an example of an immunological gain of function during adaptation to a new parasite. The three-spined stickleback, *Gasterosteus aculeatus*, is host to a specialist cestode, *Schistocephalus solidus*. Stickleback are typically marine or anadromous (migrating from the ocean into brackish estuaries to breed). During Pleistocene deglaciation (∼12,000 years ago), marine stickleback established permanent land-locked populations in newly available lakes throughout coastal north-temperate regions (24). This unique evolutionary history, with still-extant ancestral marine stickleback and thousands of replicated freshwater populations, has been widely used to study the repeatability of evolution and the genetic basis of adaptation (25–27). Most notably, the heavily armored marine fish almost always evolve reduced armor plating after establishing populations in freshwater (28, 29). But, armor defense against predators was not the only key adaptation to freshwater. When marine stickleback colonized freshwater they encountered the cestode *S. solidus*, which is exclusive to freshwater because its eggs do not hatch in brackish water (30, 31). Because they do not encounter *S. solidus* in marine habitats, the marine stickleback that colonized freshwater would have been highly susceptible to this parasite (31), and become heavily infected (32). The permanent freshwater populations subsequently evolved immune adaptations to resist *S. solidus* infection (31). For example in 1968 marine fish were experimentally introduced to a human-made quarry pond, were rapidly infected, then evolved effective resistance over the subsequent 50 years, nearly eliminating the cestode (32). This artificial population, like many natural lake populations, evolved a peritoneal-wide fibrosis response: fibrosis is induced by parasite exposure and contributes to suppressing parasite growth and viability (31). In other populations, fibrosis is actively suppressed after the initial infection, allowing the cestode to grow but avoiding pathological side-effects of fibrosis (a tolerance strategy). These differences in fibrosis response are observable in laboratory conditions, and are heritable (33). Importantly, marine stickleback, which are evolutionarily naive to *S. solidus*, lack a fibrosis response to cestodes (34, 35), though they possess the capacity for fibrosis when stimulated by artificial adjuvants. The fibrosis response, and cestode resistance more generally, thus represents an example of an immune gain-of-function when ancestral marine fish colonized freshwater and adapted to resist *S. solidus*.

However, the general mechanisms underlying fibrosis, and the evolutionary processes which led to its co-option into parasite defense in some stickleback populations, remain poorly understood (34, 36). In particular, we do not currently know what gene expression cascades lead to the initiation of peritoneal fibrosis in stickleback. Nor do we know whether this regulatory cascade is a generic response to a pro-inflammatory stimulus (for instance), or whether it is specifically a response to *S. solidus*. Finally, we do not know whether the gene regulatory response that produces fibrosis is an ancient conserved pathway shared by all stickleback populations, or whether the expression response itself is evolving to differentially regulate fibrosis in diverging stickleback populations. That is, do marine stickleback have the same gene expression responses as their freshwater relatives? Do relatively resistant and susceptible freshwater populations exhibit the same regulatory changes when they encounter *S. solidus*?

To answer such questions, we experimentally exposed lab-raised stickleback from three populations (ancestral marine, a relatively resistant lake population, and relatively tolerant lake population), to different immune challenges (alum, an adjuvant; or cestode protein extracts; and a saline control). A previous paper reported the dynamics of the fibrosis response in this experiment, showing population differences in fibrosis responses (35). To expand on that prior result, here we present the temporal regulation of gene expression in response to these different immune challenges, in the three populations. We evaluate the gene regulatory foundations of fibrosis, response to cestode proteins, and population differences in these regulatory processes. We show that the population differences in fibrosis entail changes in the same overall gene regulatory pathways, including many evolutionarily ancient processes that were repurposed during adaptation to *S. solidus*. Nevertheless, there are also population-specific responses at some genes, which highlight the capacity for immune regulatory systems to exhibit substantial change over microevolutionary time (∼10,000 generations).

## METHODS

### Experimental Design

Full details of the laboratory injection experiment can be found in Hund et al. 2022 (35). Briefly, *Gasterosteus aculeatus* were collected in June of 2018 from populations on Vancouver Island, and reproductively mature adults were used for *in vitro* breeding to generate embryos. Fish were collected from three geographically isolated and genetically divergent populations (34) Sayward Estuary (SAY) is populated with anadromous stickleback that represent a proxy for phenotypes and genotypes of the cestode-susceptible ancestral marine fish that colonized freshwater lakes on Vancouver Island (37). Importantly, this ancestor-like population has very little natural exposure to *S. solidus*. In the lab, these fish are highly susceptible to *S. solidus* and demonstrate minimal fibrosis (33, 34). Roselle Lake (RSL) stickleback have faster and stronger fibrosis response to infection (35), and consequently exhibit a lower infection prevalence in nature (7-40%; (34, 38)). The cestodes that do establish in Roselle fish tend to be smaller or even encased in a granuloma that frequently leads to parasite death (Bolnick, *pers. obs.* & Hund *unpublished data*). Because of this effective fibrosis response, we consider Roselle Lake fish to represent a resistant population, similar to the highly fibrotic Roberts Lake population studied previously (31). A second lake population, Gosling Lake (GOS) was chosen to represent a more tolerant low-fibrosis strategy. Since we began studying *S. solidus* infections in 2005, Gosling Lake stickleback typically harbored a high prevalence of *S. solidus* (50-80%), which grew rapidly to a large size without inducing discernable fibrosis. Genomic signatures of selection within fibrosis QTL suggest this population historically experienced selection favoring susceptible but tolerant genotypes that suppress the costly fibrosis response (34). We consider this population to be ‘tolerant’, with the caveat that between 2012 and 2022 the population has been undergoing rapid evolution (and declining infection rates); therefore, at the time of sampling for this study this population was likely polymorphic for fibrosis traits (39). Fish from each of the three populations were bred using standard *in vitro* fertilization methods, to generate within-population crosses. The resulting fertilized eggs were transported to the lab for rearing (40). Fish were split across two aquarium rooms at the University of Connecticut where they were reared for ∼11 months prior to the start of the injection experiment (May 2019).

At the start of the experiment, fish from each population were given intraperitoneal injections of one of three inoculants: a control injection of 20 uL of 1x phosphate-buffered saline (PBS, control), a 20uL injection of Alum (2% Alumax Phosphate, OZ Bioscience) diluted 1:1 in PBS, and a 20 uL injection of cestode protein homogenate diluted 1:1 in PBS. Alum is a common immune adjuvant which induces an innate immune response through recruitment of leukocytes (41); it had previously been demonstrated to induce peritoneal fibrosis in stickleback (Dr. Natalie Steinel, *per. comm*.). Cestode proteins were generated from *S. solidus* collected from a fourth location, Farwell Lake, which were collected in 2008, flash frozen, and stored at −80 until the time of experimentation. Use of cestode protein from a fourth population ensured that no one population of fish were challenged with native parasites to which they might be locally adapted (or which might be locally adapted to them). Full details regarding homogenate preparation are found in Hund et al. 2022. At the time of experimentation, fish were anesthetized and then injected with a randomly assigned treatment using sterile ultrafine syringes (BD 31G 8 mm). Injections were performed in the lower left peritoneal cavity, using a shallow angle parallel to the body to prevent damage to internal organs. Fish were also labeled with a small elastomer mark posterior to the neurocranium to allow for the identification of treatments (fish were housed in common tanks). Following injection, fish were allowed to recover from anesthesia before being returned to their tanks of origin. A full schematic of the experimental design can be found in **Supplementary** Figure 1.

Fish were euthanized for measurement of fibrosis and collection of tissues (pronephros and peritoneal tissue) at four time points: 1, 10, 42, and 90 days post injection. Immediately following euthanasia, fibrosis was scored following procedures described in Hund et al. 2022. Once this was complete, the pronephros or head kidneys were removed from the fish using sterile dissecting tools. Sampled tissue was suspended in RNAlater, stored at 4C overnight and then placed in −80C for long term storage. Approximately ten fish per population and treatment combination were sampled and sequenced at each the first three time points (Days 1, 10 & 42; **Supplementary Table 1**). Day 90 represented opportunistic sampling of remaining fish and had much lower and more variable sampling sizes, so it was excluded from most analyses herein with the exception of trajectory analyses.

### RNA Extraction, Sequencing and processing

RNA was extracted from head kidney tissues using an Ambion MagMax-96 RNA Isolation Kit following existing protocols (42). Extracted RNA was sent to the UT Genomic Sequencing and Analysis Facility for TagSeq Library preparation (43) and sequencing on a NovaSeq platform. All resulting reads were processed using the iRNAseq pipeline (44) which includes initial read deduplication, adaptor trimming and quality filtering. Reads were aligned to version 5 of the UGA stickleback transcriptome assembly (45–47) using bowtie2 (48).

### Gene Expression Analysis

Raw data and code for all analyses described herein can be found at GitHub (https://github.com/lfuess/InjectionMS). Prior to analyses, reads were normalized using the variance stabilizing transformation in the R package DESeq2 (49). We then filtered expression to retain only those in the upper two quartiles of relative expression (abundant mRNAs). An important caveat, as in any tissue-level transcriptomics study, is that variation in mRNA relative abundance can be attributed to several different biological processes. A given gene’s mRNA might be abundant (or rare) because it is up- or down-regulated, in the sense that chromosome unpacking and transcription factors (repressors) actively increase the rate of transcription per cell. However, one must keep in mind alternatives, such as variation in mRNA degredation rates within cells, and changes in cell populations. The latter is especially important for immune organs where cell proliferation and migration play key roles in an immune response, changing the relative abundance of different types of cells. For instance an increase in T cell numbers would increase the abundance of mRNAs expressed primarily in T cells.

For each abundant gene, we used a quasibinomial general linear model (GLM) to test whether its read count (proportional to total read depth) depended on population (SAY, GOS. RSL), treatment (PBS vs Alum; or PBS vs cestode protein in a second GLM), and sample time point. The GLMs included all pairwise and three-way interactions. A significant main effect of treatment (or, treatment*time) without any interactions with population (population*treatment or population*treatment*time) implies that there is a transcriptomic response to injection which is shared by the three populations. An interaction between treatment*time would highlight genes whose abundance responded to injection contents, but where this effect changed over time. A treatment*population interaction indicates genes that show population differences in response to injection (and treatment*population*time indicates genes where population differences in injection response shift through time). For the purposes of this analysis we do not discuss the effects of population, time, or population*time, as these model terms focus on potentially constitutive population differences (or, effects of age) unrelated to experimental treatment and therefore not germane to the questions we seek to answer. We use ‘differential expression’ (DE) to describe genes whose mRNA abundance differs between treatments (e.g., alum vs saline), keeping in mind the caveat above that multiple processes affect mRNA abundance.

Using a stringent cutoff for significance (α = 0.01), we expect that false discovery alone (type II error) would result in about 130 genes (1% of the ∼13,000 being tested with the GLMs) reaching our threshold for statistically significantly differentially expressed (DE) for a given effect. In addition to the usual adjusted threshold approach, we tested whether there are more differentially expressed (DE) genes than could be explained by type II error alone: Using a binomial proportion test we checked whether the count of DE genes exceeds the 1% null expectation for each model term (treatment, treatment*time, treatment*population, and treatment*population*time).

To evaluate similarities between DE response to alum, and to cestode protein, we wished to know how many genes exhibit DE to both treatments (relative to saline controls). If the two sets of DE genes are independent, with proportions x and y responding to alum and protein respectively, a null expectation is that a proportion x*y will be significant for both. We used a binomial proportion test to compare the observed overlap in DE genes, against this null expectation. Furthermore we tested for a correlation between effect sizes (log-2 fold change, L2FC) for both treatments using a Pearson correlation. Correlations were run on two sets of data: 1) just genes which were significantly differentially expressed in one or both models for the factor of interest, and 2) all genes tested by both models.

To further investigate population differences in transcriptomic responses, we split our samples by population and ran additional binomial GLMs with the terms treatment, time, and treatment*time, for alum and cestode population independently. We then used correlative analyses (Pearson correlations) to test for cross-population congruence in responses of genes which were significantly differentially expressed in one or more populations using possible pairwise comparisons.

Next, we considered the role of fibrosis in driving observed patterns of differential expression using a third quasibinomial GLM which tested whether read count depended on fibrosis score or the pairwise interaction of fibrosis score and population. This model was run for the entire dataset (days 1-42, control, alum and cestode protein treated), but focused on the general correlation of fibrosis with gene expression and did not consider differences in fibrosis as a result of time or treatment. Fibrosis scores from Hund et al. 2022 were used, which are the average fibrosis score across both sides of the fish (injected and uninjected). Significantly differentially expressed genes were compared across the three models for downstream interpretation. For all analyses, our discussion specifically focuses on those genes identified as putative immune genes based on updated annotations associated with version 5 of the UGA stickleback transcriptome assembly (45–47).

Finally, we conducted a trajectory analysis to evaluate the temporal progression of transcriptomic responses to the immune challenge in each population, as introduced by Torres et al. (50). To do so we generated DAPC (discriminant analysis of principal components) axis scores for each individual, using all combinations of population, treatment, and time point as grouping variables. Analyses were conducted on a reduced set of genes which were identified as significant (α = 0.01) for at least one of our model terms. We then calculated the mean DAPC score for each time point, in each population. Control fish represent the unactivated resting state, a substitute for time zero gene expression profiles. We then calculated the multivariate vectors connecting each time point, to the successive time point, for a given population. For each population we can then represent the ‘arc’ of the immune response as a series of end-to-end vectors in DAPC space, connecting successive time points. In principle, these arcs should return to their resting state if full recovery is achieved (50), providing a visual representation of immune ‘resilience’.

## RESULTS

Differential expression modeling using quasibinomial GLMs revealed targeted impacts of alum and cestode protein injections on the transcriptome composition of the stickleback head kidney (**Figure 1; Supplemental File 1**). Main effects of alum injection were most significant, inducing changes in 1,365 genes out of 13,033 tested (∼10.5%, far greater than our null expectation; binomial proportion test; *p* < 0.001). Nearly all of these genes (1,347 total) were significantly differentially expressed as a result of the main effect of alum only (no interaction terms significant), indicating strong effects of alum independent of population or timing. Of these genes, 56 were significant following Bonferroni correction at an FDR of 10% (7 immune functioning, 3 related to T cells: gpnmb, lck, and BCL11B). The interaction of alum treatment effects and time induced changes in 195 genes, significantly greater than our null expectation (binomial proportion test; *p* < 0.001). Two of these genes were significant following Bonferroni correction at an FDR of 10% (1 immune: pim1). This interaction indicates that there is a time-course of genes that are transiently expressed (or, repressed) in response to treatment, similarly for all populations. In contrast, alum by population interaction (57 genes) and three-way interactions (47 genes) both induced less differential expression than expected under false discovery rate expectations alone (binomial proportion test; *p* < 0.001). Furthermore, none of these genes were significant following a Bonferroni correction, so we cannot confidently infer any population-level differences in transcriptome response to alum at the level of individual genes.

**Figure 1:**
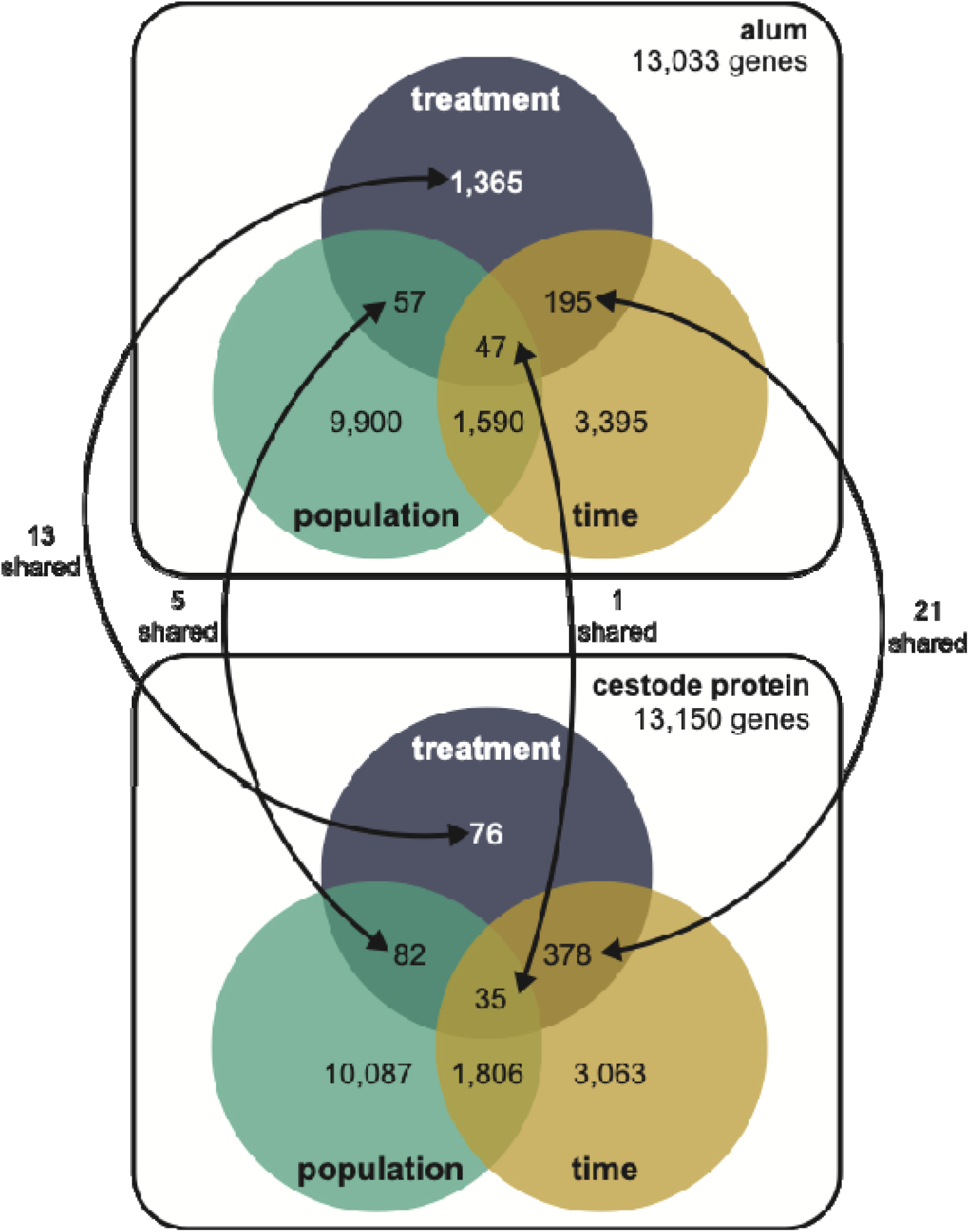
Conceptual summary of differential expression results for alum and cestode protein models. Total number of genes tested for each are displayed underneath the model title. Numbers displayed represent total differentially expressed genes for each main effect or interaction effect (where circles overlap). Arrows between the two diagrams indicate shared significant genes across both models. For the purposes of this manuscript we focused on effects (main and interactive) associated with treatment only.

In contrast to alum, main effects of cestode protein treatment were significantly less than expected, with only 76 genes significant for treatment main effects out of 13,150 tested (0.57%; binomial proportion test; *p* < 0.001). All but one of these genes were significantly differentially expressed as the result of the main effect of cestode protein only (no interaction terms significant). However, we did observe a large number of genes which changed in response to cestode protein over time; 378 genes were significantly dependent on the interaction of treatment and time, greater than false discovery alone could yield (test of equal proportions; *p* < 0.001). This suggests that cestode protein treatment induces a dynamic response over time, rather than the persistent effects that characterized responses to alum. All other terms, including treatment by population interaction (82 genes), and three-way interaction (35 genes) induced significantly less differential expression than predicted under null expectations (test of equal proportions; *p* < 0.001). No genes differentially expressed in our cestode protein model were significant after Bonferroni correction (FDR of 10%).

Finally, our GLM analysis of expression associations with fibrosis yeilded 2,888 genes whose expression was dependent on fibrosis (regardless of treatment and time), and 389 genes which were significantly dependent on the interaction of fibrosis and population (**Supplemental File 1**). Both of these exceeded null expectations (binomial proportion test*; p*<0.001). Of those genes with significant main effect of fibrosis, 147 also were significant for the interaction of fibrosis and population. Finally, 658 genes were significantly dependent on fibrosis score following Bonferroni correction at an FDR of 10%, while only one was significant post correction for the interaction of fibrosis and population. These results point to a strong transcriptomic signature of the general fibrosis response, regardless of treatment type, time, or, to some extent, population.

### Shared Responses to Alum and Cestode Protein Injection

When comparing differential expression responses to alum and cestode protein across main and interaction effects we see limited patterns of conserved responses (**Figure 1**). Only 13 genes were differentially expressed (uncorrected p<0.01) in response to both treatments, marginally-significantly greater than null expectation of ∼8 genes (test of equal proportions; *p* = 0.095). Responses to alum and cestode protein were largely positively correlated, both when considering only significantly differentially expressed genes (**Figure 2a**), and when considering all genes tested in both analyses (**Supplementary** Figure 2a). This positive correlation confirms that there is a shared basis to the transcriptomic response to alum, and to cestode protein. This shared basis of alum and protein respose is more obvious when we consider similarities in treatment*time and treatment*population effects. A total of 21 genes (∼6 expected) exhibit significant DE treatment*time interaction for alum and protein treatments, more than false discovery expectations (test of equal proportions; *p* < 0.001). Effect sizes for alum and protein were also positively correlated whether we considered just the subset of differentially expressed genes (**Figure 2b**) and when considering all genes (**Supplementary** Figure 2b). Finally, five genes exhibited treatment*population effects for both alum and protein injections (more than the <1 null expectation), again demonstrating strong congruence across treatments. Notably, only a few of these shared genes across all effects are related to immunological processes, with no clear trends in shared pathways (**Supplementary** Figure 3).

**Figure 2:**
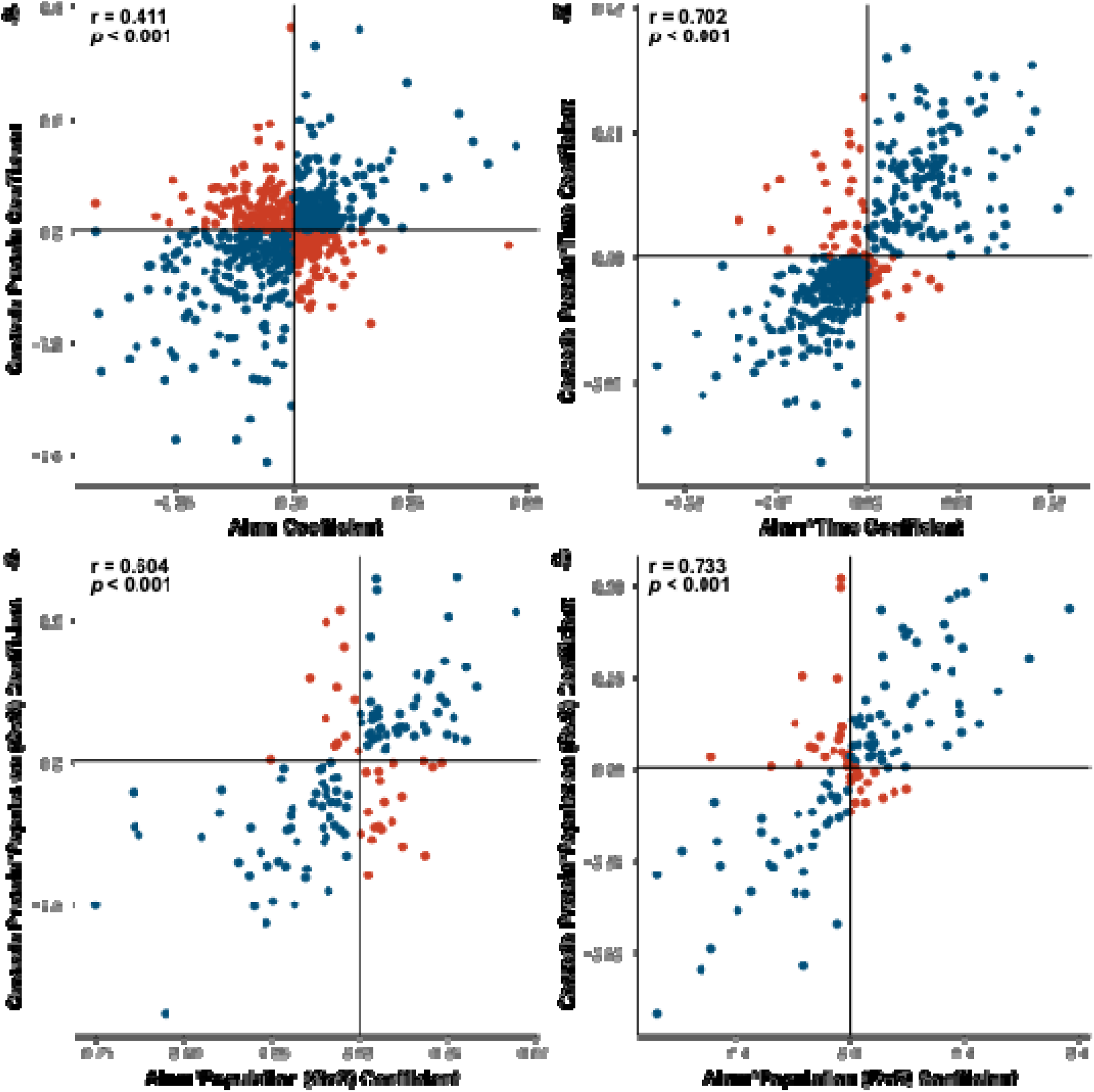
Responses to treatment are largely congruent across alum and cestode protein. Scatterplots display relationship between responses to **a)** treatment main effects, **b)** treatment by time effects and **c-d)** treatment by population effects across alum and cestode protein treatments. All points represented a gene which is significantly differentially expressed in at least one of the two models (alum or cestode protein) Points are colored based on the relationship of the coefficients for each model, wherein blue dots indicate congruence across alum and cestode protein effects and red points indicate divergence. Pearson correlation results are displayed for each comparison.

Genes which exhibit DE in response to treatment also exhibit DE in interaction terms, which provides strong evidence this DE is biologically real and not type II error. In particular we noticed significant overlap between genes which were significant for a main effect of alum and those which changed over time in response to cestode protein. A total of 54 genes were differentially expressed as a result of both alum main effects, and cestode protein*time interactions, significantly greater than the expected 40 genes (test of equal proportions; *p* = 0.0230). This observation shows that the same sets of genes react persistently to alum, and transiently to cestode protein. Eight of these genes have immunological functions, all but one of which were consistently upregulated as a result of alum. In contrast, all these genes in cestode protein treated fish started upregulated on day 1 (compared to PBS) but ended up downregulated by day 42 (**Figure 3**). This suggests that within this core set of genes responding to both treatments there is a compensatory down-regulation unique to worm protein responses. Furthermore, seven out of eight of these genes were significantly correlated to fibrosis, two of which were also differentially correlated to fibrosis across populations (klf2b and mrc1b; **Figure 4**). All of these genes were also significantly differentially expressed as a result of population in the fibrosis model, and all were significantly higher expressed in RSL compared to GOS and SAY (Tukey post-hoc; **Supplementary Table 2**; Supplementary Figure 4). Furthermore, for those genes with fibrosis*population effects, neither was significantly positively associated with fibrosis in RSL. Specifically, klf2b was significantly positively correlated with fibrosis in GOS and SAY only (population-specific quasibinomial GLM; **Supplementary Table 3**) and mrc1b was significantly positively associated with fibrosis in GOS only (population-specific quasibinomial GLM; **Supplementary Table 3**).

**Figure 3:**
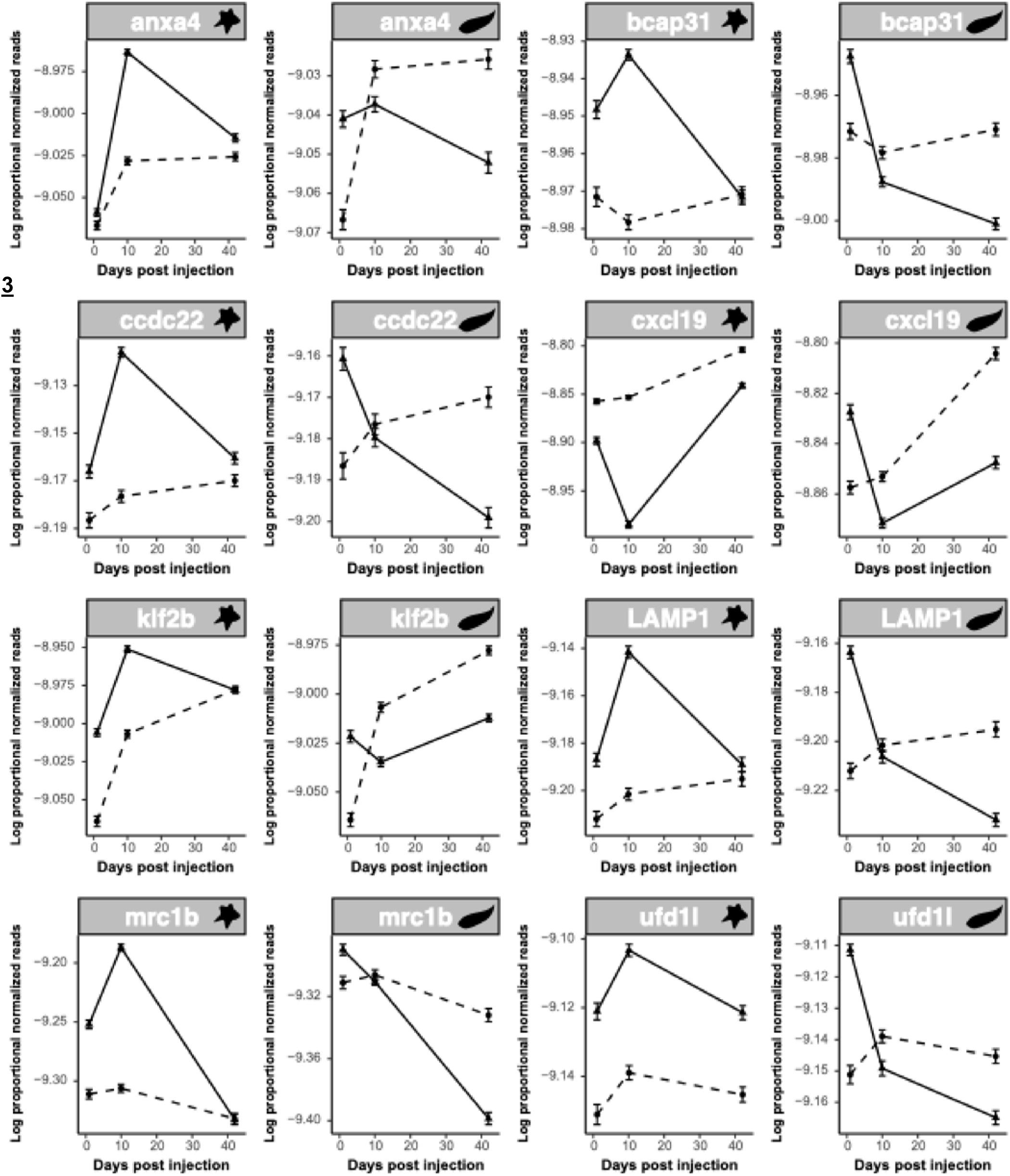
Responses to cestode protein are more dynamic over time when compared to alum. Line plots display proportional normalized read count values of putative immune genes which were differentially expressed as a result of both alum main effects and cestode protein by time effects. Plots show trajectories of treatment and control groups over time. Plots are paired with alum response on the left (indicated with star icon) and cestode protein response on the right (indicated with worm icon). Dotted lines indicate control values whereas solid lines indicate treatment values. As there were no significant population effects, lines are shown for all fish within a treatment combined across populations.

**Figure 4:**
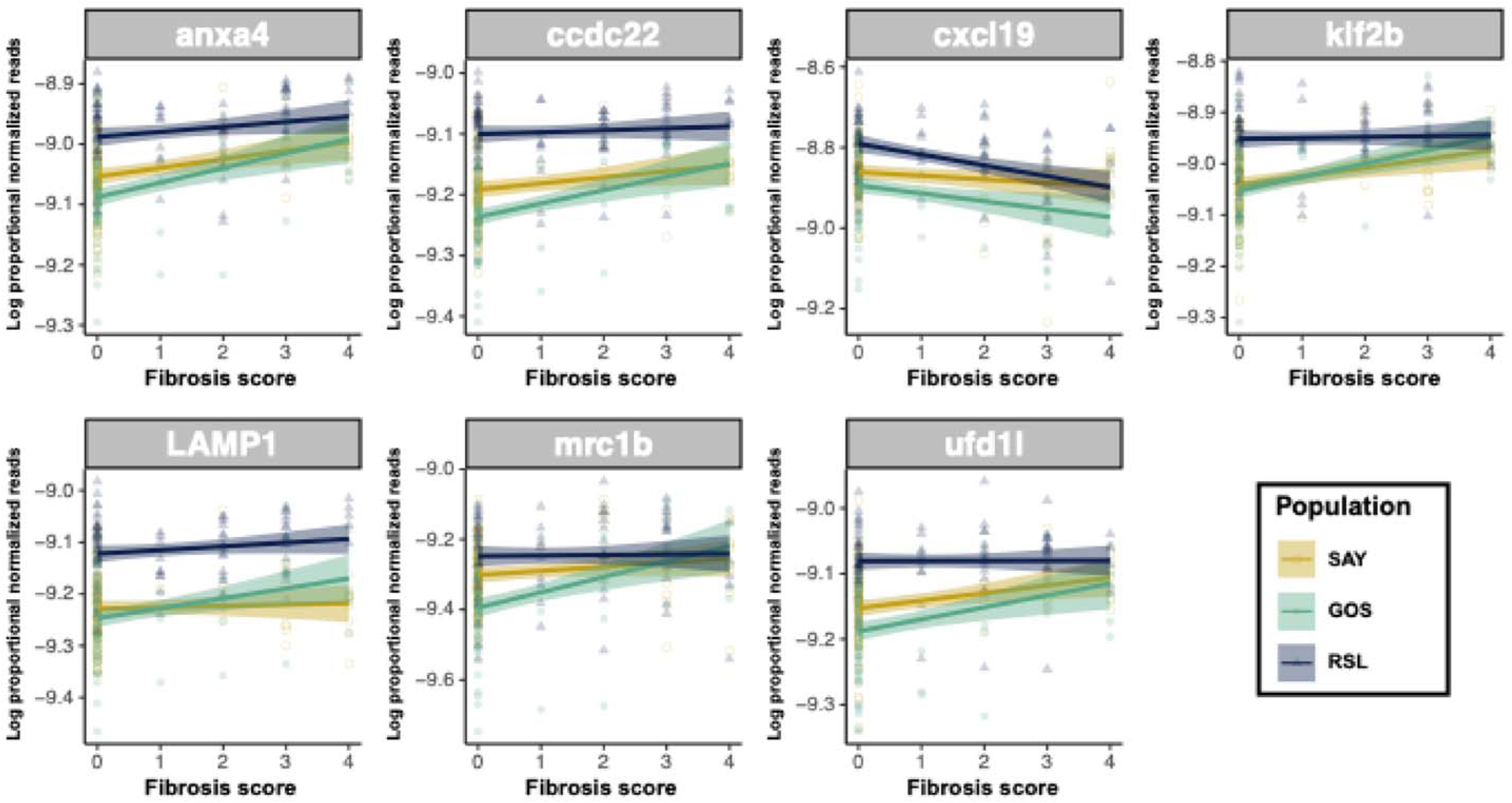
Genes which were more dynamic over time in response to cestode protein, but also responsive to alum, are associated with fibrosis. Scatter plot display associations between proportional normalized read count values and fibrosis scores for genes which were differentially expressed as a result of both alum main effects and cestode protein by time effects. Points and lines are colored by population; lines represent population-specific linear models with 95% confidence intervals shaded.

### Unique Responses to Alum and Cestode Protein Injection

Of the genes responsive to the main or interactive effects of alum, 110 were identified as putative immune genes, 46 of which (nearly half) were related to T cell function or antigen presentation. Nearly all these genes were uniquely responsive to alum treatment or alum by time interactions. A total of 35 genes which may be related to T cell function were significantly differentially expressed dependent on alum main effects, with complex associations between alum treatment and activation/function of genes associated with a wide variety of T cell subtypes (**Figure 5a**). Of those, 25 were also associated with either fibrosis or the interaction of fibrosis and population; the effect sizes of genes responses to alum treatment were positively correlated with the genes’ responses to fibrosis (pearson correlation; *p* < 0.001, r = 0.666; **Figure 5a**). This tells us that the transcriptional basis of fibrosis is driven by T cell response that is similarly responsive to alum. All but three of these genes (nr4a.1, rc3h1b, and skap1) were also significantly differentially expressed as a result of the main effect of population in the fibrosis model (**Supplementary Table 2).** Furthermore, 18 of these were significantly differentially expressed in RSL compared to GOS and SAY, all but three of which were constitutively expressed higher in RSL (Tukey post-hoc; **Supplementary Table 2).** Furthermore, of the five genes with a fibrosis by population interaction effect, none were significantly associated with fibrosis in RSL, and four of five were significantly associated in both GOS and SAY (population-specific quasibinomial GLM; **Supplementary Table 3**). From these population effects we infer that T cell responses associated with fibrosis are population-specific.

**Figure 5:**
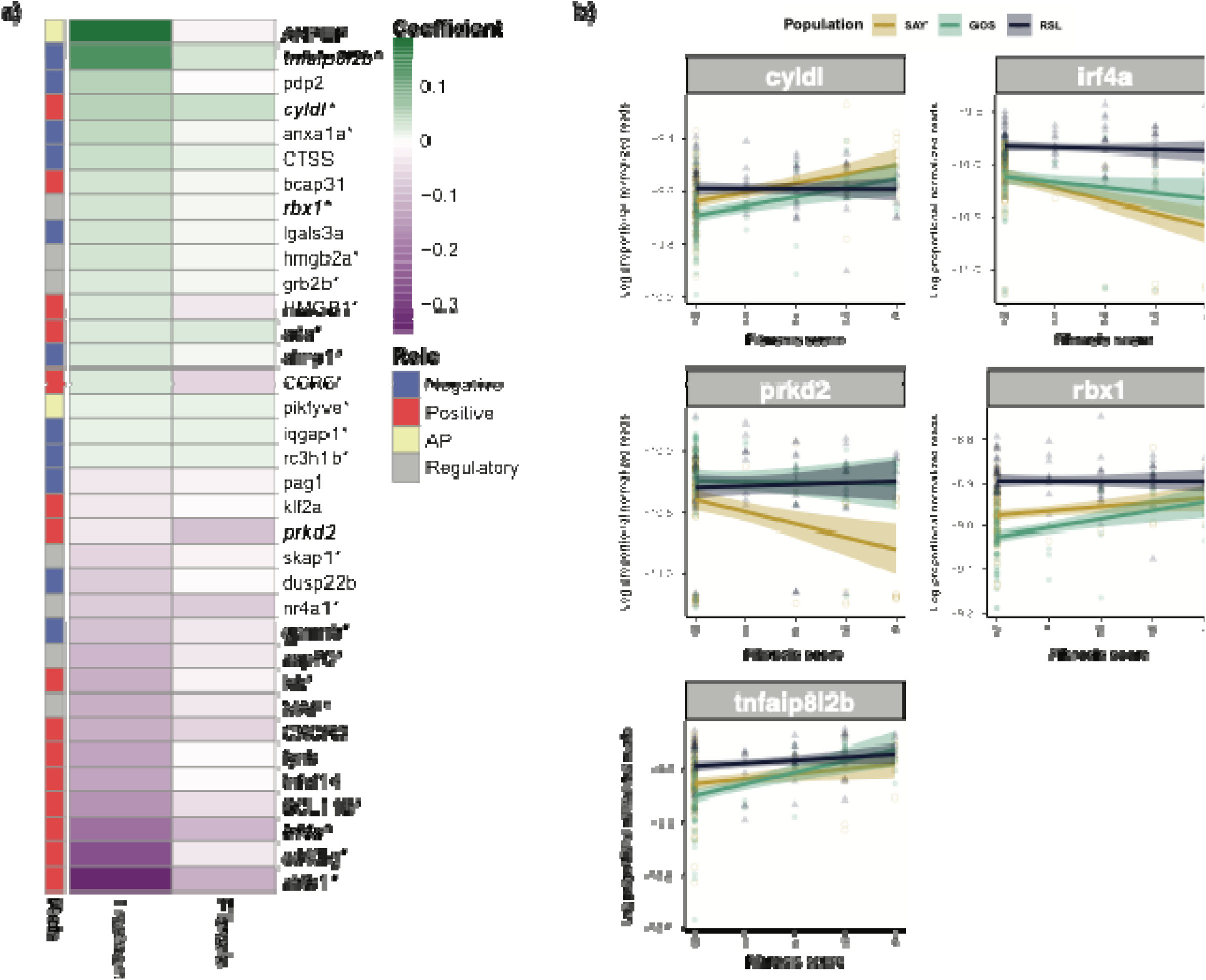
Summary of alum main effects and fibrosis effects on the subset of T cell related genes which are uniquely responsive to alum. **a)** heatmap displaying coefficient values for alum and fibrosis main effects for our unique alum genes which are associated with T cell function; The annotation bar summarizes the relationship of each gene to T cell function (POS-positive effect on T cell functioning, NEG-negative or suppressive effect on cell functioning, AP-antigen presentation, REG-contributes to the general regulation of cell function in what could be positive and/or negative ways). * indicates significance for fibrosis main effect. Bolded & italicized gene names indicate those which also had a significant effect of fibrosis by population. **b)** Scatter plot displaying associations between proportional normalized read count values and fibrosis scores for T cell genes which were differentially associated with fibrosis across genotype. Points and lines are colored by population; lines represent population-specific linear models with 95% confidence intervals shaded. Plots include both alum and cestode protein data. In summary, genes associated with T cell activity show largely congruent responses to alum and associations with fibrosis, but those with opulation specific responses are more associated with fibrosis in GOS and SAY fish, but constitutively highly expressed in RSL fish.

Several T cell associated genes were also differentially expressed in response to alum over time, all of which begin upregulated at day 1 (relative to PBS) but end up downregulated by day 42 (**Figure 6**). This up-then down-regulation is indicative of an activiation stage followed by compensatory repression, similar to those noted previously. Only two of these nine genes were also impacted by fibrosis, both negatively. Finally, of the 102 alum responsive genes with population interaction effects, only four have roles in immunity, three of which are related to T cell function. Each of these is differentially responsive to treatment in GOS compared to RSL and SAY fish (**Supplementary** Figure 4). None of these overlap with fibrosis.

**Figure 6:**
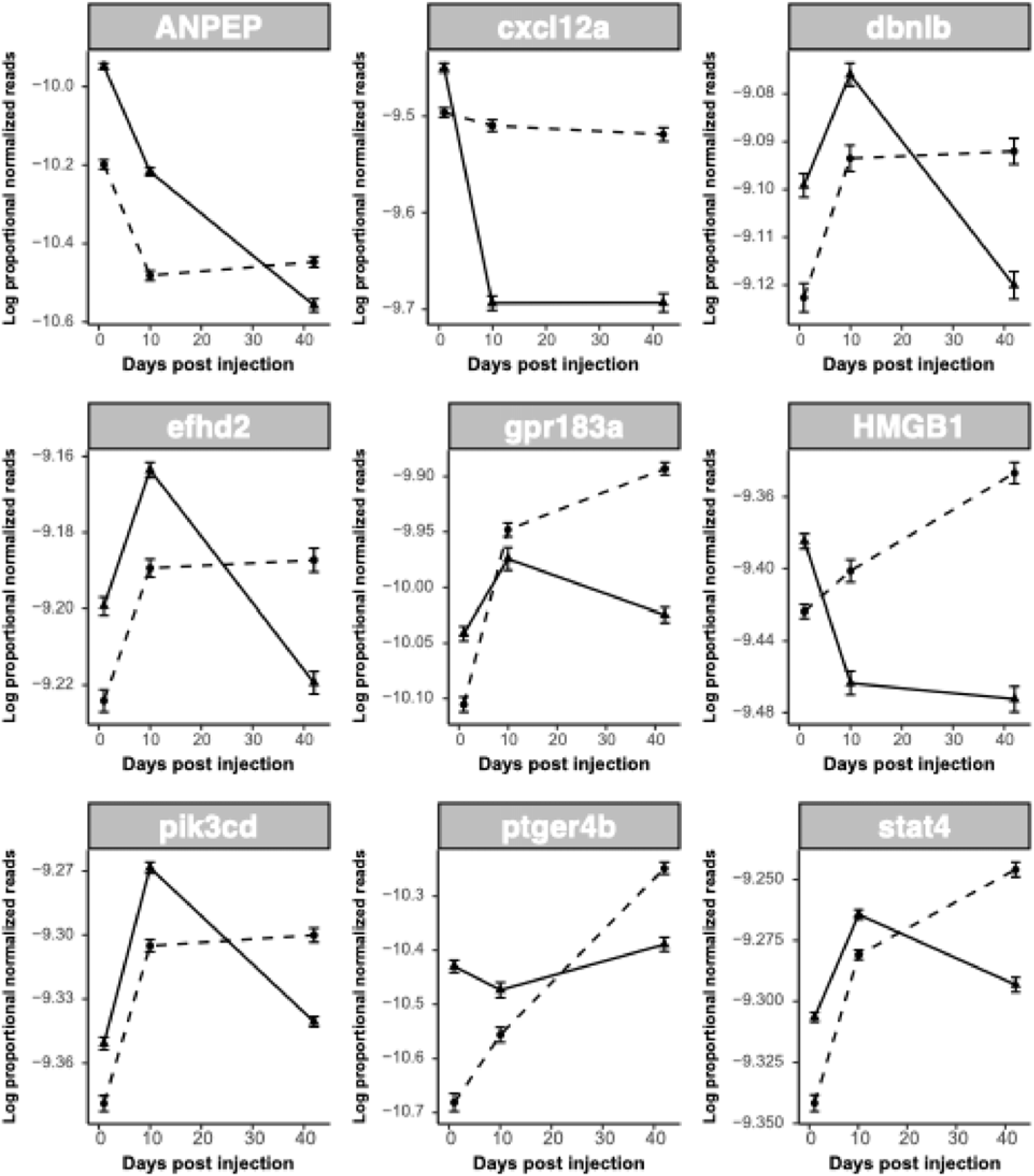
Some T cell genes which are responsive to alum display dynamic patterns of regulation; most start upregulated and end up suppressed. Line plots display proportional normalized read count values of uniquely alum-responsive T cell genes which were differentially expressed in response to alum over time. Plots show trajectories of treatment and control groups over time. Dotted lines indicate control values whereas solid lines indicate alum treatment values. As there were no significant population effects lines are shown for all fish within a treatment combined across populations.

When considering unique responses to cestode protein, a clear pattern of enrichment of genes involved in activating and regulating the function of key immune transcription factor, NF-kB, is observed. Thirteen of the 39 identified immunological genes affected by cestode protein main or interactive effects are related to NF-kB functioning (9 unique to cestode protein). Of the unique genes, seven are significantly differentially expressed in response to cestode protein over time; all but one of these genes begin upregulated at day 1 (compared to PBS) and end up downregulated by day 42 (**Figure 7**). Again, this suggests a cycle of gene activation then repression. Only three of these overlap with fibrosis; all three were also significant for population and were expressed higher in RSL compared to GOS/SAY (Tukey post-hoc; **Supplementary Table 2).** Furthermore, mul1a and nfkbie are negatively associated with fibrosis regardless of population while commd7 is positively associated with fibrosis in GOS and SAY only (population-specific quasibinomial GLM; **Supplementary Table 3**).

**Figure 7:**
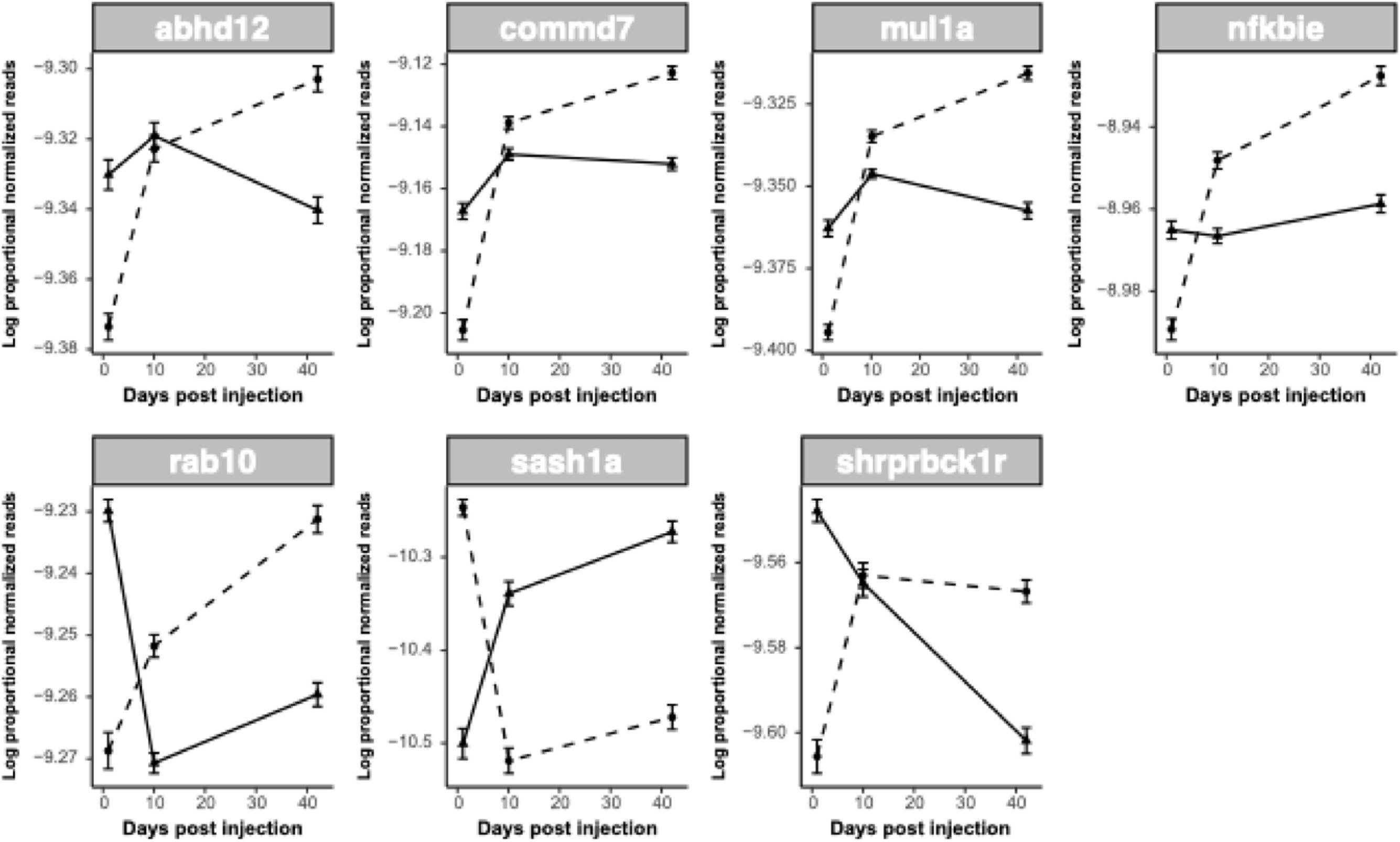
Many NF-kB genes which are responsive to cestode protein display dynamic patterns of regulation; most start upregulated and end up suppressed. Line plots display proportional normalized read count values of uniquely cestode protein-responsive NF-kB genes which were differentially expressed in response to cestode protein over time. Plots show trajectories of treatment and control groups over time. Dotted lines indicate control values whereas solid lines indicate cestode protein treatment values. As there were no significant population effects lines are shown for all fish within a treatment combined across populations.

Related to NF-kB functioning, we also see strong enrichment of inflammatory-related genes, some of which overlap with our NF-kB genes. Twelve of our 39 immune-related cestode protein differentially expressed genes are associated with inflammatory processes (7 unique to cestode protein). Of the unique cestode protein inflammation genes, five are affected by the interaction of population and cestode protein treatment effects, or the three-way interaction of all effects (treatment, population, time; **Figure 8**). All five are divergent in GOS fish compared to RSL and SAY fish. Only one of these, dusp10, is also associated with fibrosis (negatively).

**Figure 8:**
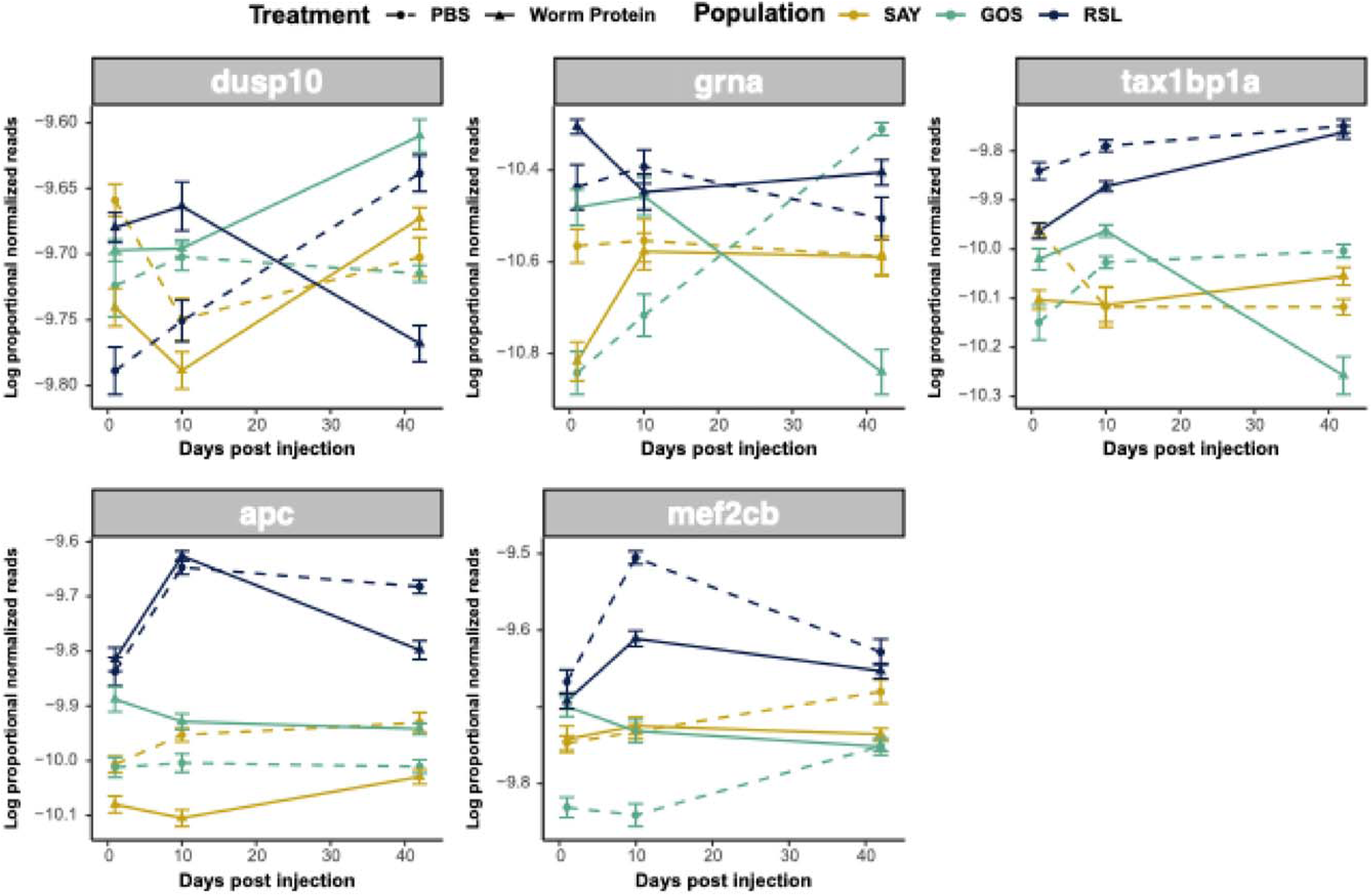
Inflammatory responses to cestode protein are population specific. Line plots display proportional normalized read count values of uniquely cestode protein-responsive inflammation genes which were differentially expressed in response to cestode protein across population. Plots show trajectories of treatment and control groups over time. Lines are colored based on population. Dotted lines indicate control values whereas solid lines indicate cestode protein values. Genes are sorted based on effects-the top row displays genes with significant three-way interactions (treatment by population by time) whereas the bottom row shows genes with treatment by population effects.

Finally, putative fibrosis regulator gene spi1b was also significantly differentially expressed in the cestode protein group only and was differentially associated with fibrosis across populations (**Figure 9**). Spi1b was initially upregulated in response to cestode protein but downregulated by day 42. Notably, the gene was significantly differentially expressed across populations (highest in RSL; Tukey post-hoc; **Supplementary Table 2**) and strongly positively associated with fibrosis in GOS and SAY fish only (population-specific quasibinomial GLM; **Supplementary Table 3**).

**Figure 9:**
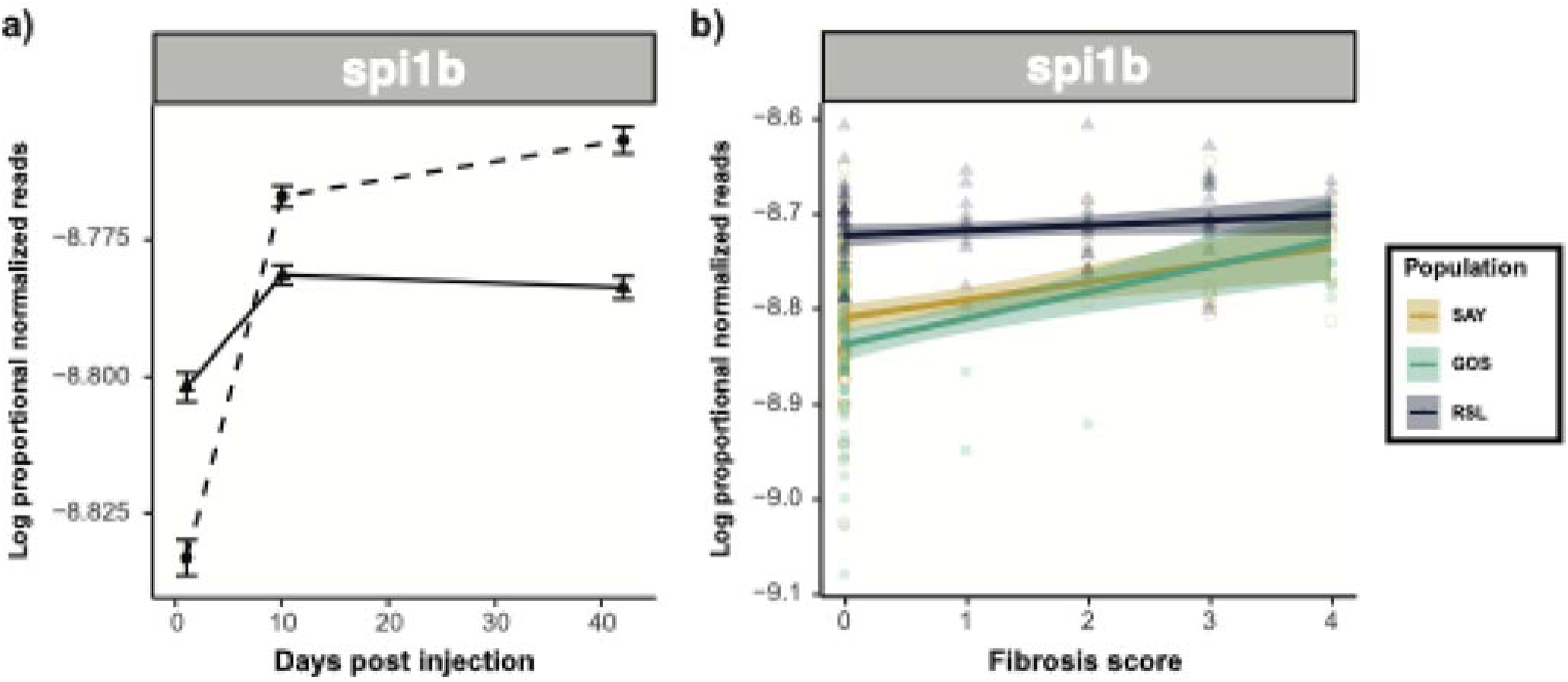
Summary of differential expression of putative stickleback fibrosis regulator gene spi1b in response to cestode protein treatment and its effect on fibrosis. *Spi1b* is dynamically responsive to cestode protein treatment (initially upregulated, but eventually downregulated) and positively associated with fibrosis in GOS and SAY only. In RSL fish it is constitutively highly expressed. **a)** Line plots displaying proportional normalized read count values trajectories of treatment and control groups over time. Dotted lines indicate control values whereas solid lines indicate cestode protein values. As there were no significant population effects lines are shown for all fish within a treatment combined across populations. **b)** Scatter plot displaying associations between proportional normalized read count values and fibrosis scores. Points and lines are colored by population; lines represent population-specific linear models with 95% confidence intervals shaded.

### Population Specific Responses

Multivariate analyses of the transcriptome can reveal broader coordinated changes spanning many genes. One multivariate approach is to compare the entire vector of differential expression (DE) effect sizes, for all genes, between populations. This transcriptome-wide approach can reveal broader scale differences in immune responses, than would be seen with individual genes (**Supplemental File 2**). Correlative analyses comparing population-specific responses to alum and cestode protein across populations confirmed general conservation of responses to both alum and cestode protein (**Table 1**), among populations. That is, in general if a gene is up- (or, down-) regulated in response to alum (or cestode protein) in SAY fish, it is likely to change in a similar direction and magnitude in GOS or RSL lake stickleback. The exception is when we compare the two lake populations’ responses to cestode protein (GOS vs RSL): for this one contrast, the correlation in DE values is negative rather than positive. Genes that are up-regulated in GOS tend to be down-regulated in ROS and vice versa. Genes with divergent cestode protein responses across GOS and RSL included antiviral gene mxb, complement gene c7, and inflammatory genes AREL1, cdo1, and dpf2; **Supplementary** Figure 5).

**Table 1:**
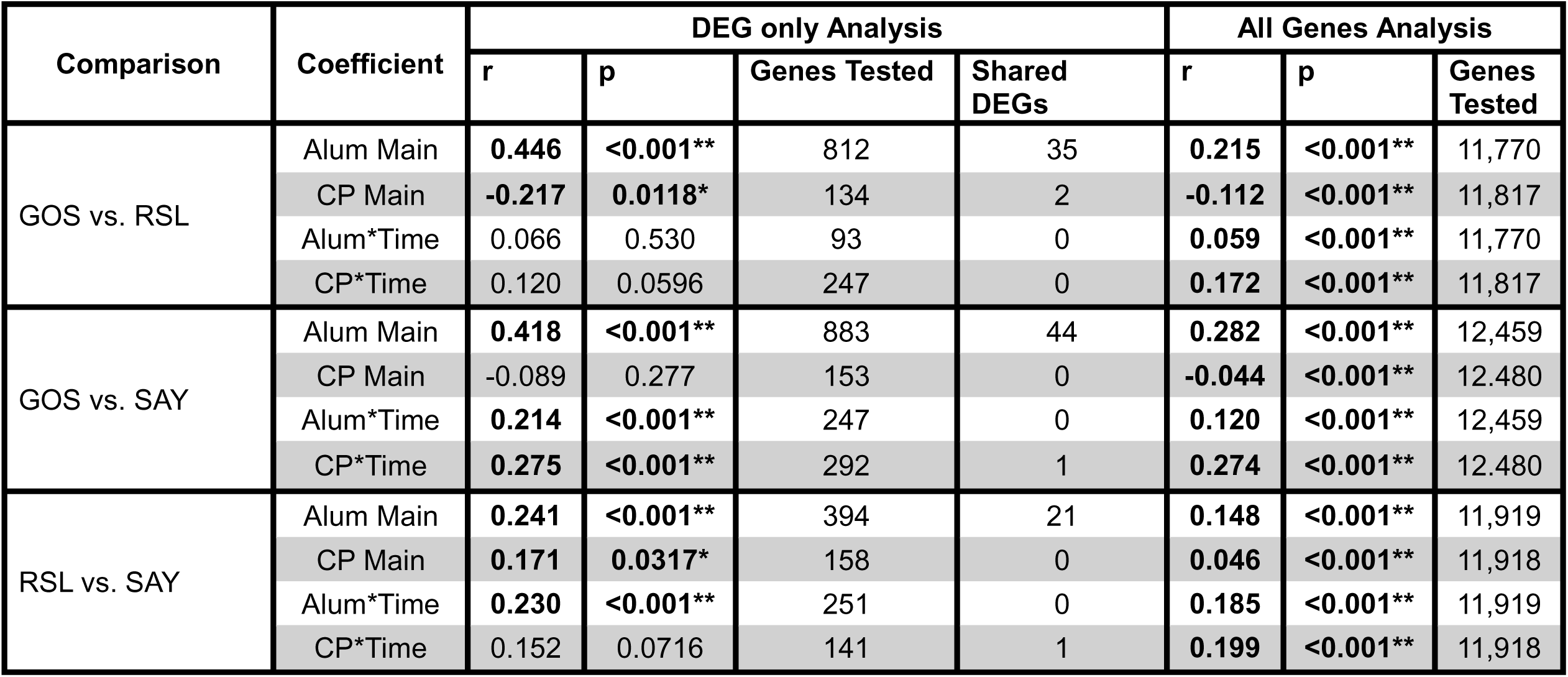
Pearson correlation results comparing treatment and treatment by time coefficients across population-specific models for both alum and cestode protein. Correlations were run only on genes that were tested in both population models (i.e., in the top two quartiles of expression for both populations). Two sets of correlations were run, one on only genes that were differentially expressed in one or both of the populations being compared, and one using all genes that fit the criteria. The number of genes tested in both analyses is noted, as are the total numbers of shared differentially expressed genes.

| Comparison | Coefficient | DEG only Analysis |  |  |  | All Genes Analysis |  |  |
| --- | --- | --- | --- | --- | --- | --- | --- | --- |
|  |  | r | p | Genes Tested | Shared DEGs | r | p | Genes Tested |
| GOS vs. RSL | Alum Main | **0.446** | **<0.001\*\*** | 812 | 35 | **0.215** | **<0.001\*\*** | 11,770 |
|  | CP Main | **-0.217** | **0.0118\*** | 134 | 2 | **-0.112** | **<0.001\*\*** | 11,817 |
|  | Alum*Time | 0.066 | 0.530 | 93 | 0 | **0.059** | **<0.001\*\*** | 11,770 |
|  | CP*Time | 0.120 | 0.0596 | 247 | 0 | **0.172** | **<0.001\*\*** | 11,817 |
| GOS vs. SAY | Alum Main | **0.418** | **<0.001\*\*** | 883 | 44 | **0.282** | **<0.001\*\*** | 12,459 |
|  | CP Main | -0.089 | 0.277 | 153 | 0 | **-0.044** | **<0.001\*\*** | 12,480 |
|  | Alum*Time | **0.214** | **<0.001\*\*** | 247 | 0 | **0.120** | **<0.001\*\*** | 12,459 |
|  | CP*Time | **0.275** | **<0.001\*\*** | 292 | 1 | **0.274** | **<0.001\*\*** | 12,480 |
| RSL vs. SAY | Alum Main | **0.241** | **<0.001\*\*** | 394 | 21 | **0.148** | **<0.001\*\*** | 11,919 |
|  | CP Main | **0.171** | **0.0317\*** | 158 | 0 | **0.046** | **<0.001\*\*** | 11,918 |
|  | Alum*Time | **0.230** | **<0.001\*\*** | 251 | 0 | **0.185** | **<0.001\*\*** | 11,919 |
|  | CP*Time | 0.152 | 0.0716 | 141 | 1 | **0.199** | **<0.001\*\*** | 11,918 |

Another multivariate approach developed by (50) uses ordination to summarize variation in gene expression among individuals and track the temporal progression of transcriptomic changes. The initial direction of multivariate change after infection (or injection) indicates the early set of genes activated to drive an immune response. Later vectors reveal sets of genes driving down-regulation and recovery, which could either lead to a new resting state (e.g., chronic inflammation) or return to the original resting state. To what extent do the stickleback populations show parallel or divergent vectors? Trajectory analyses reveal clear between-population differences in response to treatment, which again were most notable when considering cestode protein. Trajectories across the first 42 days of the experiment were largely similar across both populations and treatment types (**Figure 10**). Furthermore, no return to PBS state was observed in any group. However, when including data from day 90, we observed differences in trajectory patterns across our three populations and treatment types (**Supplementary** Figure 6). From day 42 to day 90 RSL fish only show some movement back towards the PBS baseline, with stronger movement in response to cestode protein compared to alum. In contrast, both SAY and GOS continue to move away from the PBS marker, with the most pronounced movement way observed in SAY response to alum and GOS response to cestode protein. This result is consistent with the previously published observation that RSL fish partly recover from fibrosis, whereas GOS and SAY do not. Thus, the trajectory analyses mostly indicate parallel responses early after injection, and divergence mostly in the recovery phase.

**Figure 10:**
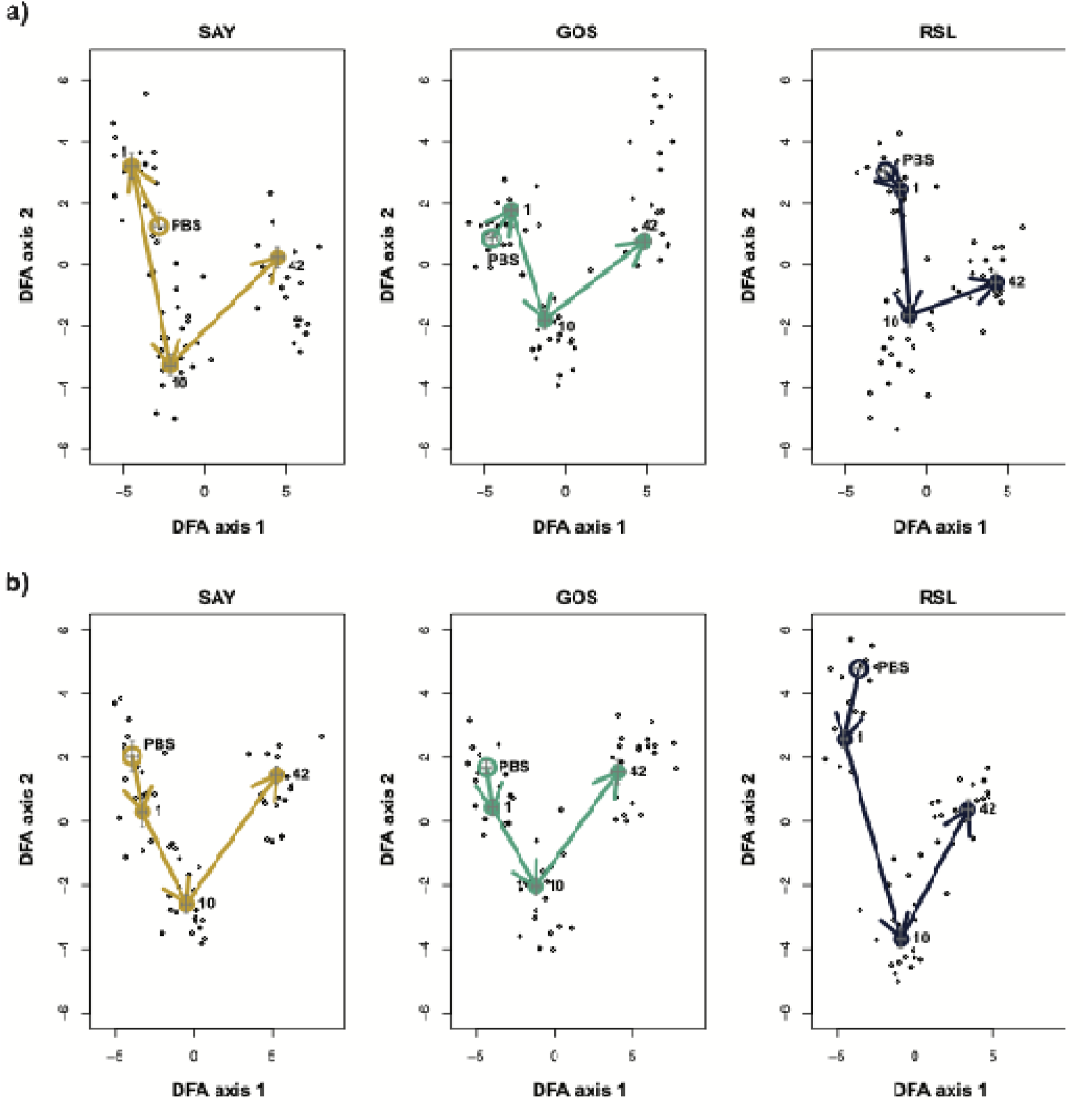
Trajectory plot of response to **a)** alum and **b)** cestode protein for each population independently based on genes which were significant for treatment or any interaction term including treatment across both alum and cp (2235 total). Closed circles indicate centroids for treated fish at each time point, whereas open circle indicates centroid for control fish. Arrows indicate trajectory between time points. Crosses indicate relative spread of data points at each time point. Axes are not standardized across plots for simplified visualization. Plots show relatively similar trajectories through day 42.

## DISCUSSION

Parasites exert immense selective pressure on their hosts and have been widely considered a key driver of host diversification, evolutionary novelty, etc (51, 52). While robust evidence suggests that these patterns extend to evolution of the immune system (13–15), many gaps still exist in our understanding of the mechanisms controlling evolutionary origins of defense responses to novel parasites. Here we present an empirical approach leveraging transcriptomic data to characterize defense responses that evolved in response to a novel parasite, and the mechanisms which have allowed a general physiological response (fibrosis) to be co-opted for parasite defense. By comparing responses to a general and parasite specific stimuli across populations we are able to highlight the roles of regulatory changes in facilitating evolution of parasite resistance. Our results reveal substantial co-option of ancient genetic pathways which remain intact, but whose expression level and timing are altered in freshwater populations that adapted to a cestode in the past 12,000 years.

### General fibrotic responses to alum are mediated by change in T cells with some patterns of self-regulation

When examining transcriptomic responses unique to alum, we see strong signatures of complex changes to T cell activity. Like any tissue-level transcriptomic study, this could reflect either increased expression per cell, or an increased abundance of a cell type (e.g., T cell proliferation or immigration) in the studied tissue. Similar changes are also strongly associated with fibrotic phenotypes. Alum adjuvant is widely used due to its immunostimulatory functions, and has been documented to have broadscale effects on host T cell activity (53, 54). Specifically, alum adjuvant is known for its impact on antigen presentation and CD4^+^T cell activation, largely through direct interactions with dendritic cells (53, 54). This activation typically is thought to bias towards a Th2-type response (55). In contrast to these previous results, our study of stickleback responses to alum reveals broad patterns of changes in genes associated with a wide variety of T cell types including CD4^+^ (i.e. aimp1 and lck (56, 57)), CD8^+^ (i.e. HMGB1 & hmgb2a (58, 59)), regulatory T cells (i.e. nr4a1, (60)), and even Th17 (i.e. pdp2, (61)). We also fail to see strong evidence of a Th2 bias; for example, anxa1a, which induces a Th1 bias (62), is upregulated in response to alum in stickleback. Our data indicate a more complex and nuanced interaction between alum and T cell activity in fish compared to mammals which is worth further investigation, especially considering broad reactivity to the adjuvant across fish species (63).

Notably, these changes in T cell gene activity have significant overlap with gene expression modeling of fibrosis; changes in gene expression in response to alum are largely congruent with effects on fibrosis score. This highlights a potential undescribed role for T cell activity in mediating fibrotic responses to alum. While fibrotic changes and granulomas have been reported in response to alum in some animal models (64, 65), understanding of this link is limited. Additionally, ample evidence has suggested a prominent role for T cell activity in fibrosis across tissue types (66, 67). The roles of T cells in regulating fibrosis are complex; Th1 cells are widely considered antifibrotic, whereas Th2 cells promote fibrosis (68). The complex patterns of T cell associated gene expression captured in our dataset in response to alum are likely reflective of these multi-faceted roles of T cells in fibrosis. It should also be noted that our data suggest that these associations between alum, T cell activity, and fibrosis are evolutionarily malleable. Notably, several alum-induced T cell genes have different associations with fibrosis depending on the fish population being examined here (**Figure 5b**).

### Defense responses to a novel parasite are heavily reliant on evolutionarily ancestral immune pathways

Although our results regarding alum are an interesting illustration of the dynamic nature of immune regulation over microevolutionary time, stickleback are not evolving to adapt to alum. Rather, they are evolving to defend against (or tolerate) diverse parasites such as *S. solidus*. The differences (and similarities) in stickleback populations’ responses to *S. solidus* protein injection can tell us more about the process of adaptation to a novel parasite. In particular, to what extent do the freshwater populations (which are frequently exposed to *S. solidus*) use deeply conserved immune regulatory pathways present in many vertebrates? Do they use pathways that are also present in the ancestral marine fish that did not co-evolve with this cestode? Or, are their responses significantly derived?

While many components of defense response to cestode protein are unique when compared to alum, they draw upon what would largely be considered ancestral, conserved immune responses. Specifically, cestode protein responses are most broadly characterized by changes in genes associated with inflammation and activity of the evolutionarily ancestral, multi-faceted immune transcription factor, NF-kB. Specifically, we see general trends of initial upregulation of NF-kB reltated genes in response to cestode protein, which revert to suppression by day 42 (**Figure 7**). NF-kB is a prominent immune regulator (69) which has frequently been co-opted into defense response to parasites across taxa (70–73). Previous evidence even suggests its role in stickleback parasite response (74). The broad scale activity of NF-kB, as well as numerous pathways for its activation likely explain its central and repeated role in parasite defenses (69).

Notably, reflective of the broad activity of NF-kB, we observe few indications of induction of specific immune responses following cestode protein treatment. Instead, we observe general changes in inflammatory-associated genes. Inflammatory responses are regulated by NF-kB activity (69) and important components of general host defense, including parasite defense (75). Inflammation is also often positively associated with, or even triggers, fibrosis (76), potentially explaining the population-specific expression patterns observed. Notably, we see tighter regulation of inflammation processes in response to cestode protein in GOS fish, mostly through increased activation of anti-inflammatory genes including dusp10 (77), tax1bp1a (78), apc (79), and mef2cb (80). Suppression of inflammatory responses following cestode protein exposure in GOS fish likely mechanistically contributes to reduced fibrotic responses.

### Further evidence supporting the role of candidate fibrosis gene, *spi1b*, in cestode defense

Finally, while general, ancestral defense responses were most common in characterizing response to cestode protein, we did also observe a response that is specific to sticklebacks’ defense against cestode protein: the induction of a key candidate fibrosis gene, spi1b. Spi1b encodes a transcription factor which has roles in fibrosis responses (81). In stickleback specifically, mounting genomic evidence suggests that evolutionary divergence in this gene and/or surrounding regulatory regions underlies differences in observed fibrosis phenotypes across population (34). CRISPR/cas9 editing of *spi1b* alters fibrosis phenotypes in stickleback, and pharmacological inhibition of the *spi1b* protein product (PU.1) leads to suppressed fibrosis response (39). Our data demonstrate that spi1b activation is restricted to response to cestode protein only, and is tightly regulated. Spi1b is quickly activated following injection of cestode protein (but not alum), but by the end of the experiment is suppressed in treated fish, further indicating the importance of regulation of potentially harmful defense responses. Finally, *spi1b* expression is also positively associated with fibrosis in two of three populations (see below for further discussion of population differences). This positive association is consistent with an experiment showing that drug inhibition of *spi1b* (by db1976) suppressed fibrosis response in lab-raised stickleback (39). On average, these two populations only initiate fibrosis in response to alum, but not cestode protein. This poses an puzzle in need of future functional characterization: why does cestode protein (but not alum) induce *Spi1b* expression in populations that fibrosis in response to alum (but not cestode protein)?

### Comparison of responses between treatments and populations reveals the importance of response reregulation in effective parasite defense

We observed few shared genes amongst main effect responses to treatments (13 genes total; only 4 immune). However, we did note significant overlap in those genes which were persistently responsive to alum (significant main effect of treatment) and those which responded transiently to cestode protein producing a treatment*time interaction (54 genes, 8 immune). These genes serve diverse immune roles, but were unified in their broad patterns; all but one were consistently up regulated in response to alum, also up regulated in response to cestode protein at day 1, but then downregulated by day 42. We see similar patterns even in unique alum and cestode protein responses. For example, nine of our alum-responsive T cell associated genes were significantly dependent on the interaction of alum and time, starting out activated in alum treated fish, but reaching a suppressed state by the end of the experiment. Likewise, inflammatory responses to cestode protein were also tightly regulated over time in some populations. These patterns are echoed by our trajectory analyses, wherein we observe potential regulatory changes or negative feedback between days 42 and 90 in RSL only (fast fibrosis; cestode protein fibrosis). Thus, both comparison of responses across treatments and populations suggest that regulation of fibrosis in response to cestode protein is essential. Longer scale experiments (and with higher temporal resolution) will be key in exploring these regulatory pathways and their roles in fibrosis recovery.

Combined, these results provide strong evidence that regulation of defense responses to novel parasites like *S. solidus* is key to immune adaptation. This is likely particularly true in the case of fibrosis, which can have significant fitness consequences for the host (34, 38, 82). Costly defense responses are theorized to be tightly regulated by negative feedback loops to mitigate potential immunopathology (17, 83, 84). Here we provide compelling empirical evidence to support these theories; our data suggest that co-option of fibrosis to defend against novel parasites co-evolved with associated increased regulation of the phenotype.

### Shared and divergent parasite responses of stickleback populations

Phenotypic measures of fibrosis responses were highly variable across populations (35); Choi et al. *in prep*). RSL fish exhibited a faster fibrosis response to alum than GOS and SAY, but were also able to recover (reduced fibrosis by day 90) whereas GOS and SAY retained high fibrosis. RSL fish also responded to cestode protein whereas GOS and SAY fish did not. These striking phenotypic differences between populations were not mirrored by the transcriptomic data presented here. Our general models detected only modest population-level differences in gene expression responses to both alum and cestode protein. However, comparative analyses of our gene expression data do shed some light on potential population divergence at the transcriptomic level which might explain strong variation in treatment response. Most notably, when we consider overlap in results from our treatment- and fibrosis-focused models, we see notable patterns in population divergence in fibrosis genes. A number of the genes that responded to injection were also related to fibrosis score, including genes with both a main effect of alum, and cestode protein*time interactions, T cell associated genes, and spi1b. Several of these genes were significantly differentially associated with fibrosis across populations (cyldl, irf4a, klf2b, mrc1b, prkd2, rbx1, spi1b, tnfaip8l2b), nearly all of which displayed similar patterns. Specifically, most of these genes were tightly associated with fibrosis score in GOS and SAY fish (slower fibrosis only to alum), but not in RSL (fast fibrosis response to both alum and cestode protein). Furthermore, RSL fish frequently had the highest levels of expression of all of these fibrosis genes (those significant for both main and interactive effects).

This suggests that patterns of faster fibrosis induction, and unique fibrotic responses to to cestode protein in RSL (35), may be due to higher constitutive expression of fibrosis-associated genes (i.e. constantly “on”). This builds upon prior findings which suggested that constitutive higher expression of immune genes may contribute to parasite resistance in other stickleback populations (85). Indeed, recent work shows that some stickleback populations are severely fibrotic even in the absence of an immune challenge, evolution having apparently driven them to an anticipatory response that is always on (Choi et al, *in prep*). Furthermore, a shift toward constitutive activation of these genes also likely contributes to observed patterns of immune regulation; constitutive expression of genes which can induce costly phenotypes likely results in selective pressure for observed patterns of enhanced regulation.

Our population comparative models further highlight potentially important population-level divergence in responses. While population-specific model responses to both alum and cestode protein were largely congruent when comparing our ancestral population (SAY) to either freshwater population (GOS or RSL), we note significant divergence in response to cestode protein between our two freshwater populations, GOS and RSL (*r* = −0.217, *p* = 0.0118). This means that genes that tend to be up-regulated in response to cestode in GOS tend to be down-regulated in the other, a remarkable reversal for populations that have diverged for only about 12,000 years. These patterns are potentially evidence of divergent evolution of parasite response strategies (34), though recent adaptive introgression into GOS may have dampened our signature (39).

## Conclusions

Herein we use transcriptomic analyses to highlight the gene expression responses underlying divergent host responses to a novel parasite, with a particular focus on a potent defense response (fibrosis). Our results provide new insights regarding both general mechanisms of fibrosis (i.e. T cells activity) and parasite-specific responses (i.e. NF-kB and inflammation). Most importantly, we leverage comparative analyses to highlight the importance of immune regulation and negative feedback mechanisms in parasite-specific defenses. These findings highlight the importance of the evolution of immune regulatory mechanisms as a step towards co-option of fibrosis into parasite defense. Furthermore, we also provide new evidence that shifts away from stimuli-based induction of fibrosis and towards constitutive expression of fibrosis-associated genes underly population specific responses. These patterns are observed only in fish populations which induce fibrosis quickly and in response to cestode protein, while other freshwater (i.e. parasite exposed) fish display divergent responses. Combined, our results suggest that evolution of costly parasite defenses involves changes in baseline expression and evolution of associated immune regulatory mechanisms to prevent self-harm associated with this effective but costly immune defense. These results provide important empirical evidence to support robust theoretical study of the evolution of defense response optima.

## ACKNOWLEDGEMENTS

This work was financially supported by an AAI Fellowship to LEF/DIB, NSF EES-2334642 to LEF, Texas State University startup to LEF, a James S. McDonnell Foundation Complexity Postdoctoral Fellowship to AKH, University of Connecticut start up funding to DIB, and NIH R01 grant 2R01AI123659-07 to DIB

## Supplementary Tables

**Supplementary Table 1:** Table of sample sizes for each population-treatment-time point combination.

| Population | Treatment | Time Point | Sample Size |
| --- | --- | --- | --- |
| GOS | Alum | 1 | 10 |
| GOS | Cestode Protein | 1 | 10 |
| GOS | PBS | 1 | 11 |
| GOS | Alum | 10 | 10 |
| GOS | Cestode Protein | 10 | 10 |
| GOS | PBS | 10 | 10 |
| GOS | Alum | 42 | 10 |
| GOS | Cestode Protein | 42 | 10 |
| GOS | PBS | 42 | 11 |
| GOS | Alum | 90 | 8 |
| GOS | Cestode Protein | 90 | 2 |
| GOS | PBS | 90 | 3 |
| RSL | Alum | 1 | 10 |
| RSL | Cestode Protein | 1 | 9 |
| RSL | PBS | 1 | 9 |
| RSL | Alum | 10 | 11 |
| RSL | Cestode Protein | 10 | 10 |
| RSL | PBS | 10 | 11 |
| RSL | Alum | 42 | 10 |
| RSL | Cestode Protein | 42 | 11 |
| RSL | PBS | 42 | 10 |
| RSL | Alum | 90 | 3 |
| RSL | Cestode Protein | 90 | 8 |
| RSL | PBS | 90 | 3 |
| SAY | Alum | 1 | 11 |
| SAY | Cestode Protein | 1 | 10 |
| SAY | PBS | 1 | 10 |
| SAY | Alum | 10 | 10 |
| SAY | Cestode Protein | 10 | 11 |
| SAY | PBS | 10 | 10 |
| SAY | Alum | 42 | 10 |
| SAY | Cestode Protein | 42 | 11 |
| SAY | PBS | 42 | 10 |
| SAY | Alum | 90 | 3 |
| SAY | Cestode Protein | 90 | 4 |
| SAY | PBS | 90 | 2 |

## Supplementary Figures

**Supplementary Figure 1:**
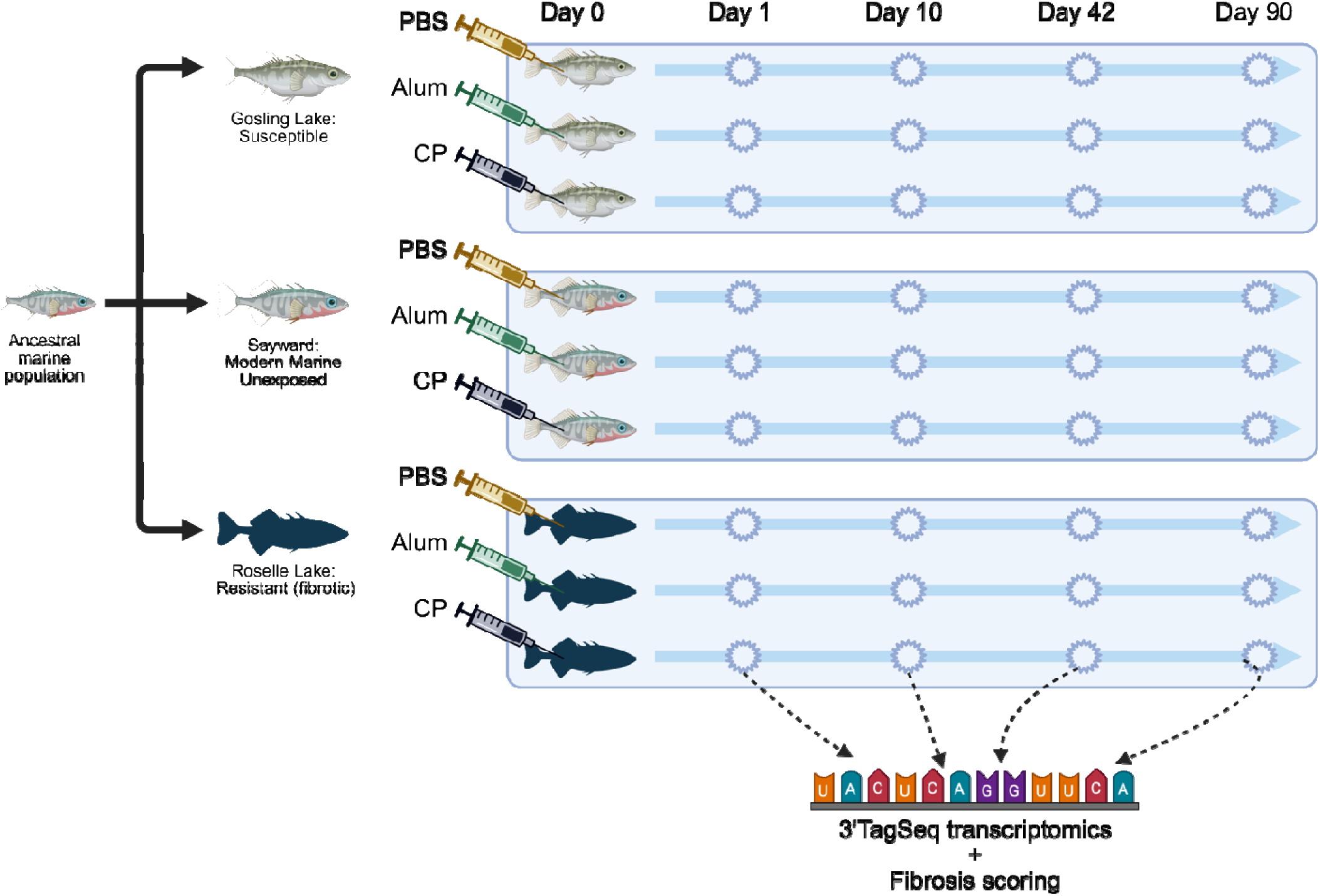
Schematic displaying experimental design setup for data collection; fish from three populations: Gosling (GOS), Sayward (SAY), and Roselle (ROS) were exposed to one of three treatments: phosphate buffered saline control (PBS), alum adjuvant (alum), or cestode protein (CP). Fish were then sampled at Days 1, 10, 42, and 90 post injection. Fibrosis was scored and head kidneys were removed for 3’ TagSeq transcriptomics.

**Supplementary Figure 2:**
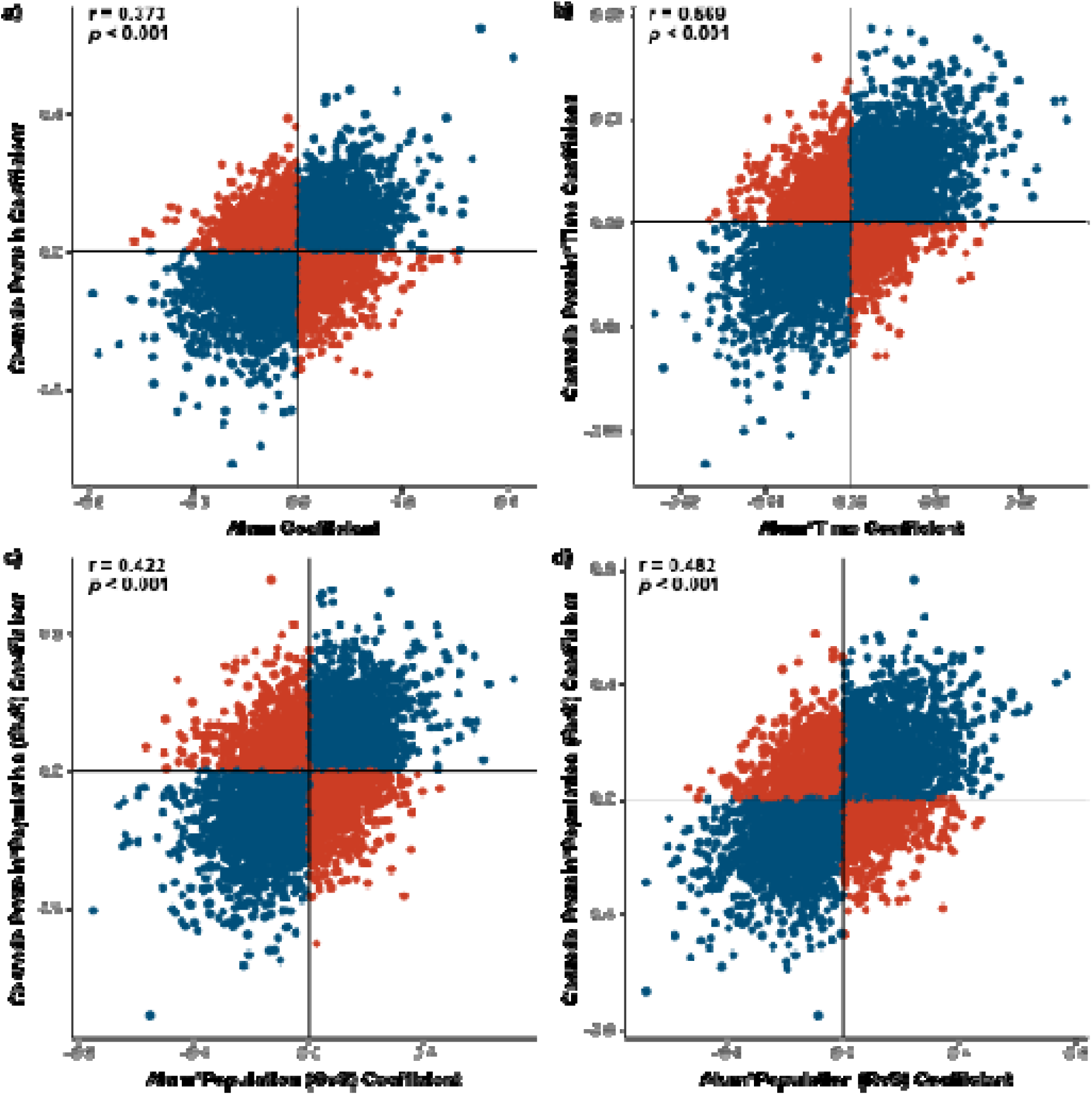
Scatterplots displaying relationship between responses to **a)** treatment main effects, **b)** treatment by time effects and **c-d)** treatment by population effects across alum and cestode protein treatments. Graphs show all genes which were tested in both models. Points are colored based on the relationship of the coefficients for each model, wherein blue dots indicate congruence across alum and cestode protein effects and red points indicate divergence. Pearson correlation results are displayed for each comparison.

**Supplementary Figure 3:**
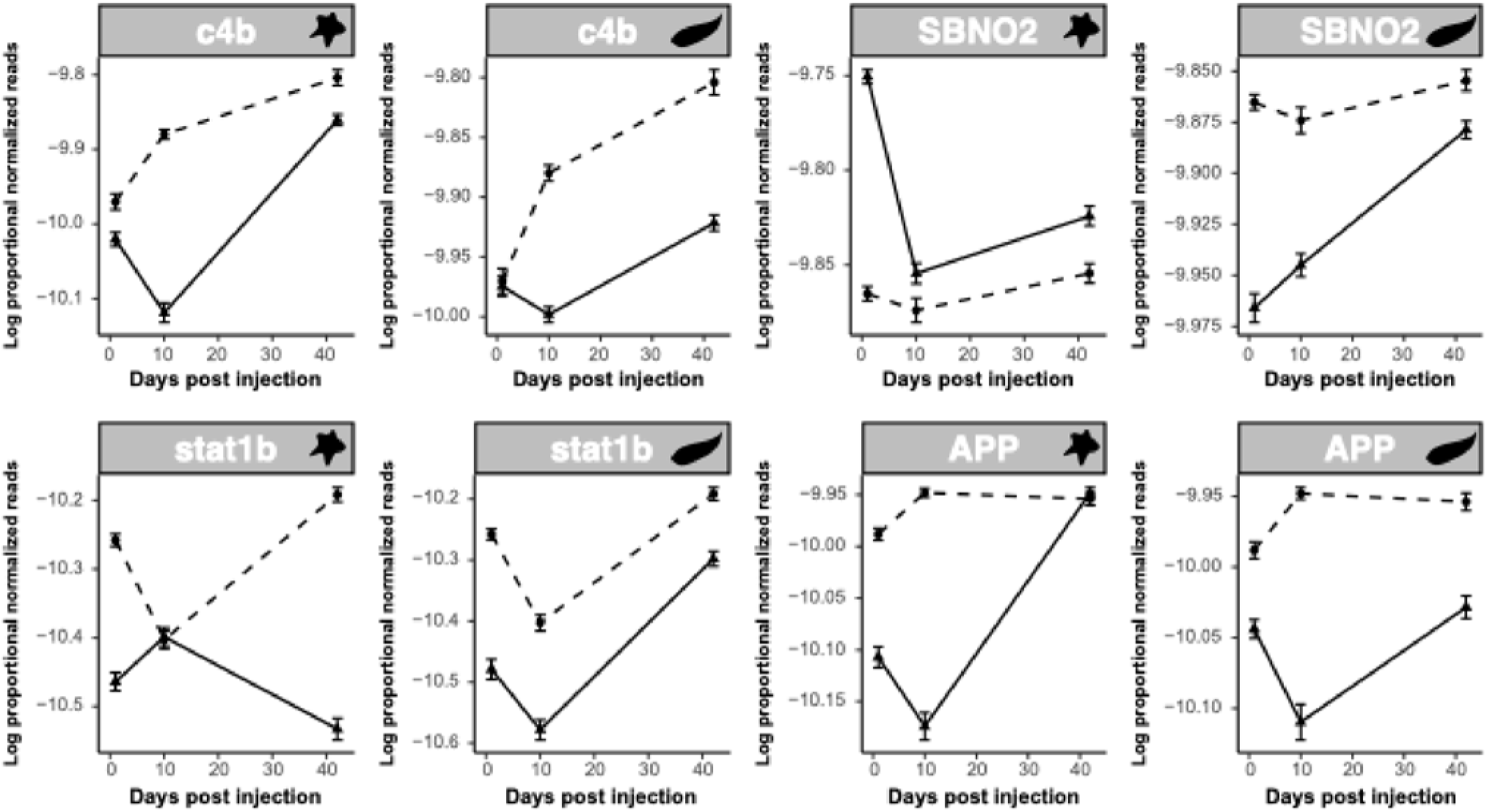
Line plots displaying proportional normalized read count values of genes of interest in treatment and control groups over time. Plots are paired with alum response on the left (indicated with star icon) and cestode protein response on the right (indicated with worm icon). Dotted lines indicate control values whereas solid lines indicate treatment values. As there were no significant population effects lines are shown for all fish within a treatment combined across populations.

**Supplementary Figure 4:**
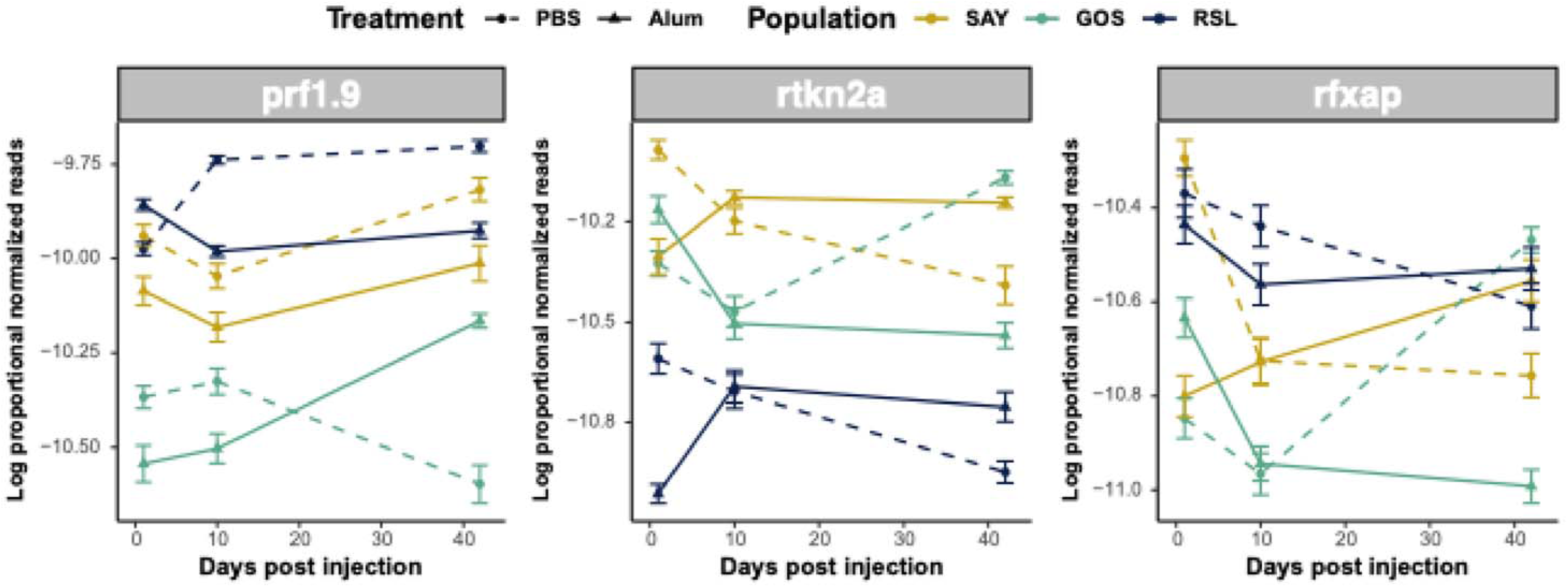
Line plots displaying proportional normalized read count values of uniquely alum-responsive T cell genes which were differentially expressed in response to alum over time and across genotypes. Plots show trajectories of treatment and control groups over time. Lines are colored based on genotypes. Dotted lines indicate control values whereas solid lines indicate cestode protein values.

**Supplementary Figure 5:**
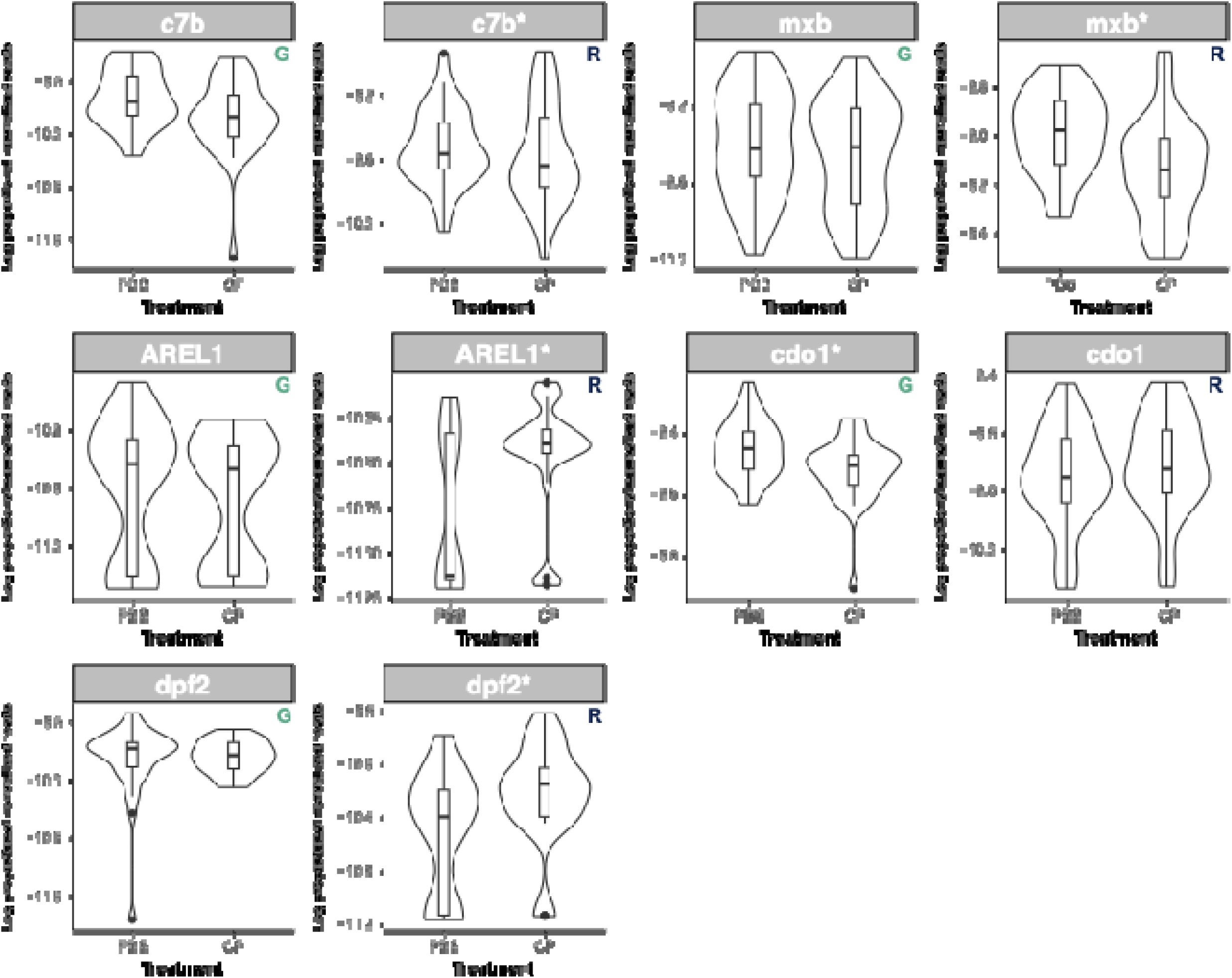
Violin plot with inlaid box and whisker plot for putative immune genes which displayed opposing responses to cestode protein in the population specific modules. * Indicates significance of the treatment term in the model. Plots are paired with GOS on the left and RSL on the right, indicated by the small colored letter in the upper righthand corner of each plot.

**Supplementary Figure 5:**
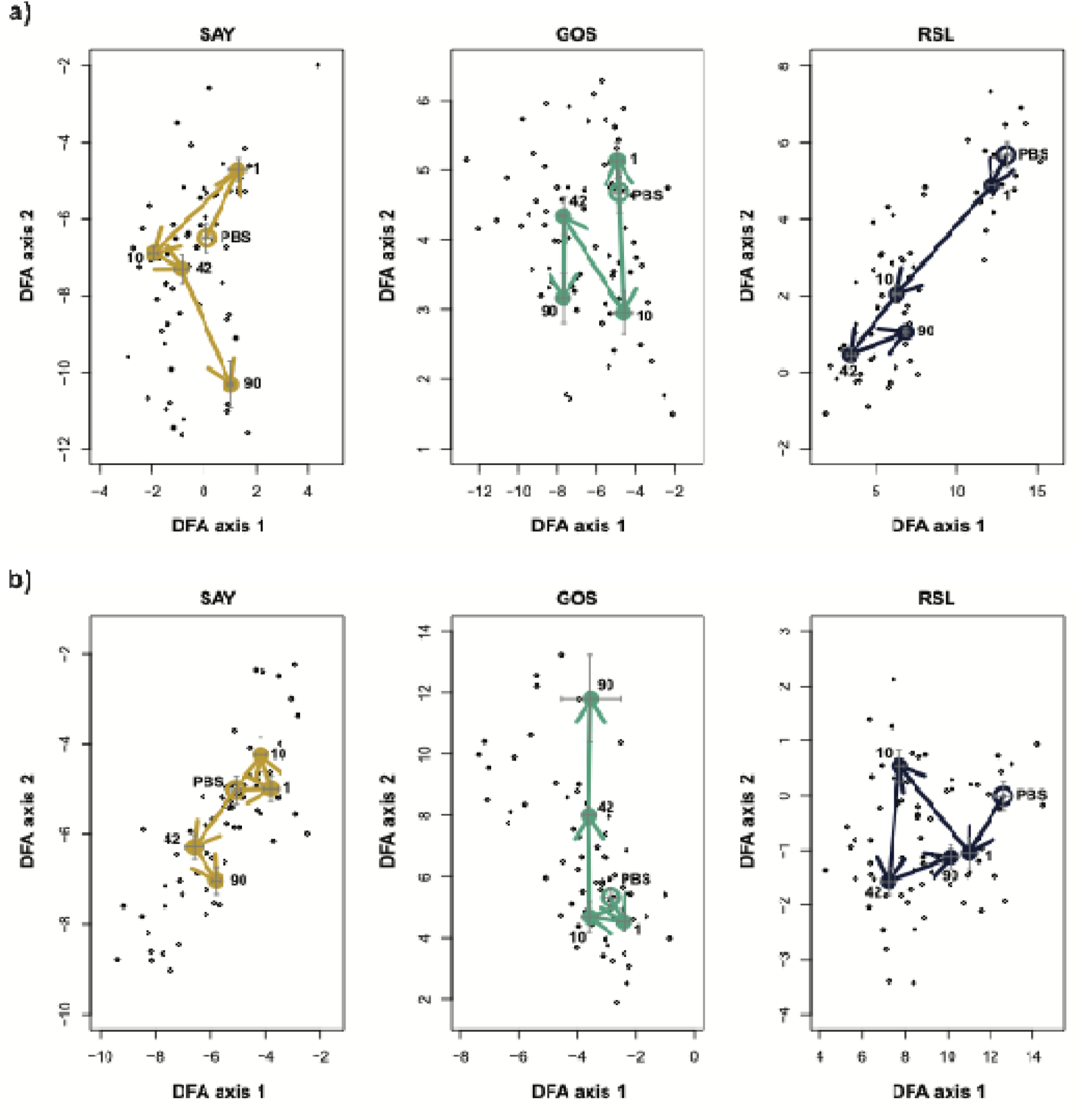
Trajectory plot of response to **a)** alum and **b)** cestode protein for each population independently based on genes which were significant for treatment or any interaction term including treatment across both alum and cp (2235 total). Closed circles indicate centroids for treated fish at each time point, whereas open circle indicates centroid for control fish. Arrows indicate trajectory between time points. Crosses indicate relative spread of data points at each time point. Axes are not standardized across plots for simplified visualization.

## Notes

### Competing Interest Statement

The authors have declared no competing interest.

