## Supplementary figures and images for "Changes in gene regulation are associated with the evolution of resistance to a novel parasite"

### Supplementary Figure 1

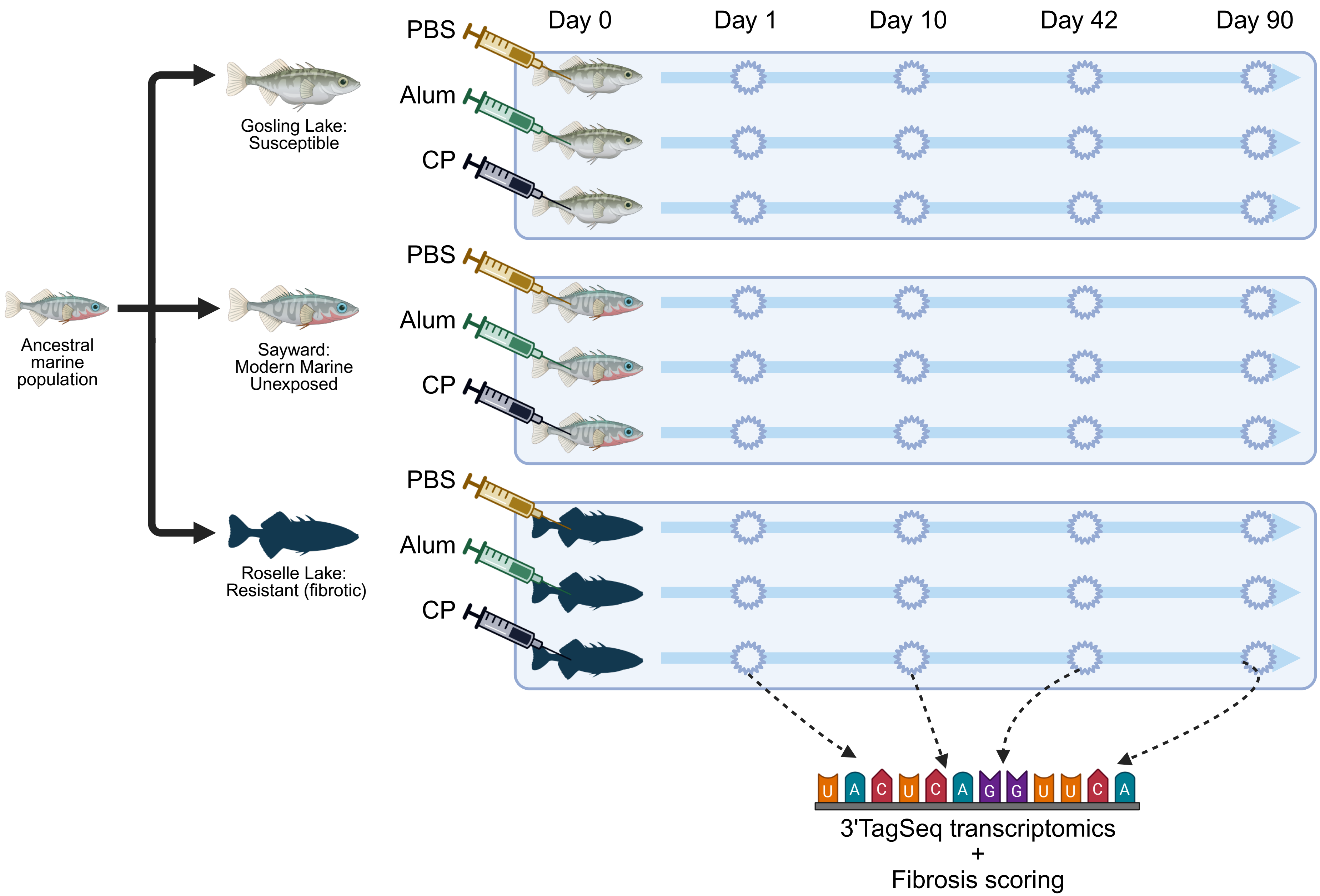

### Supplementary Figure 2

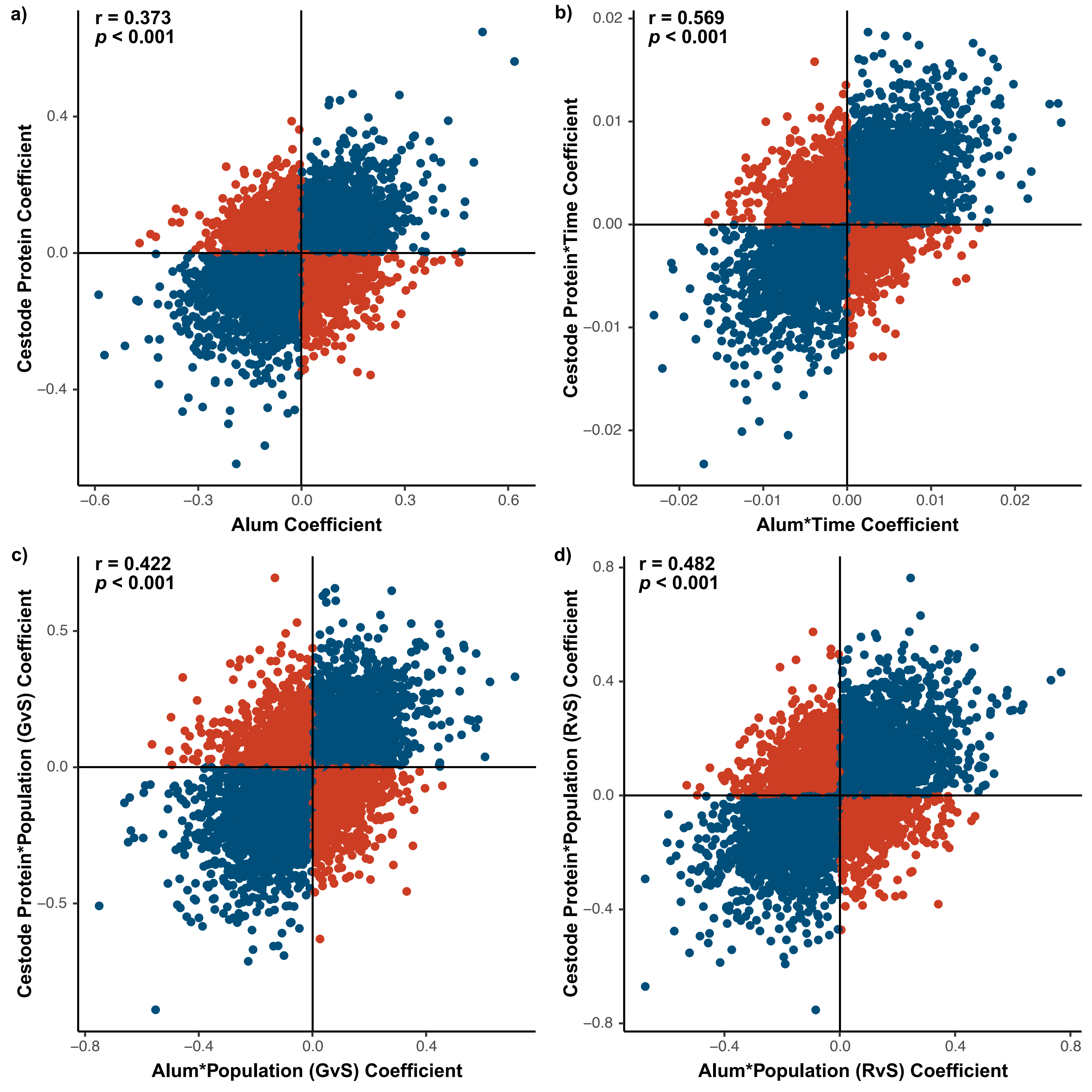

### Supplementary Figure 3

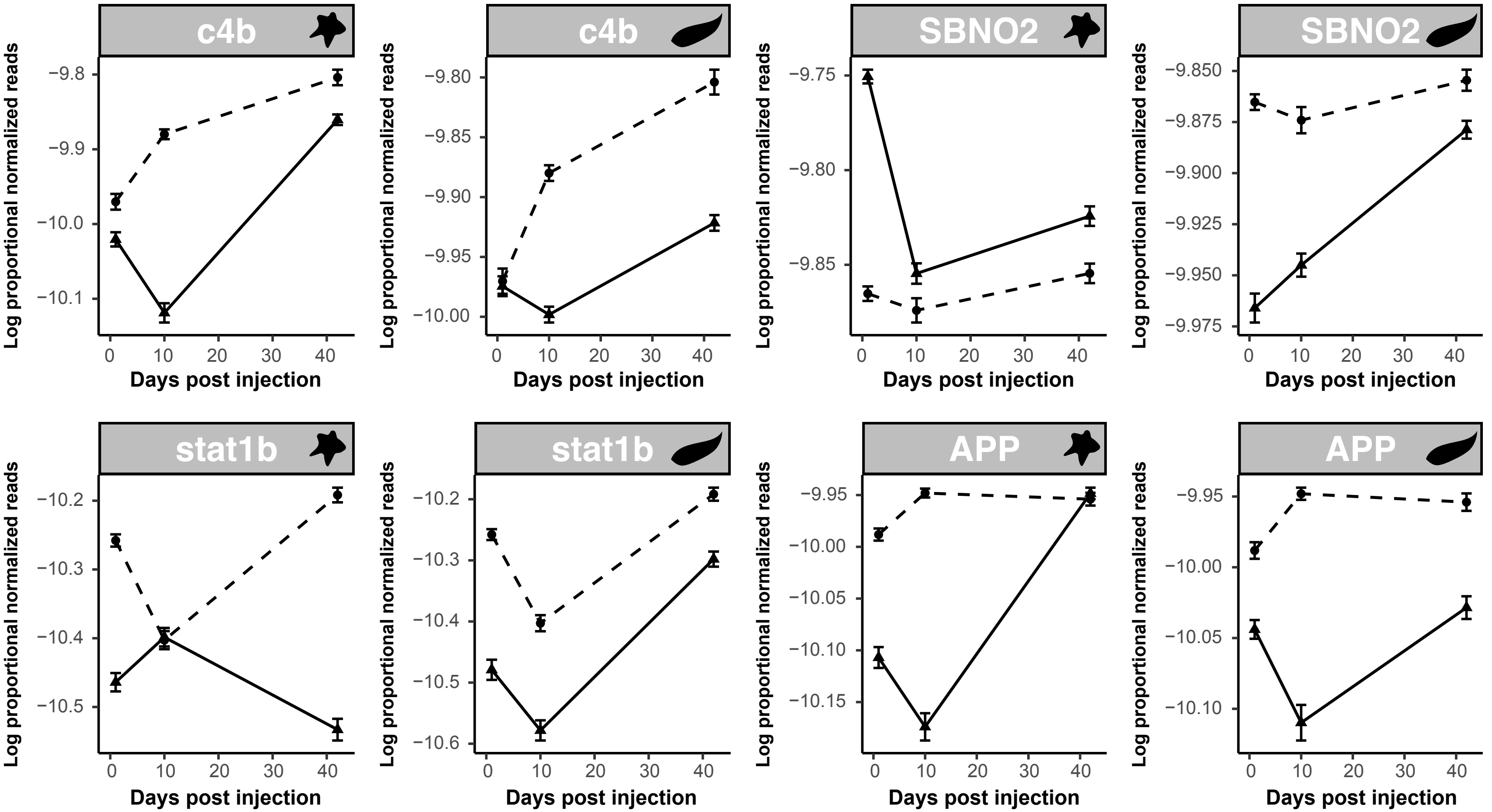

### Supplementary Figure 4

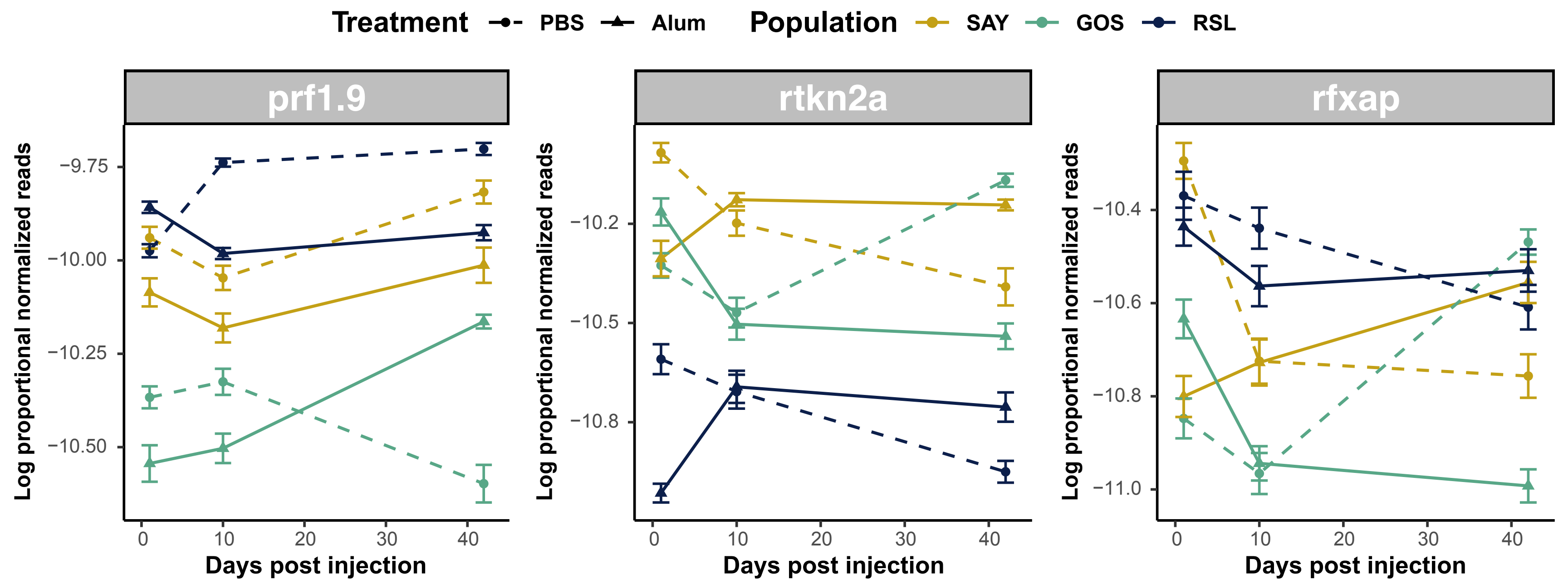

### Supplementary Figure 5

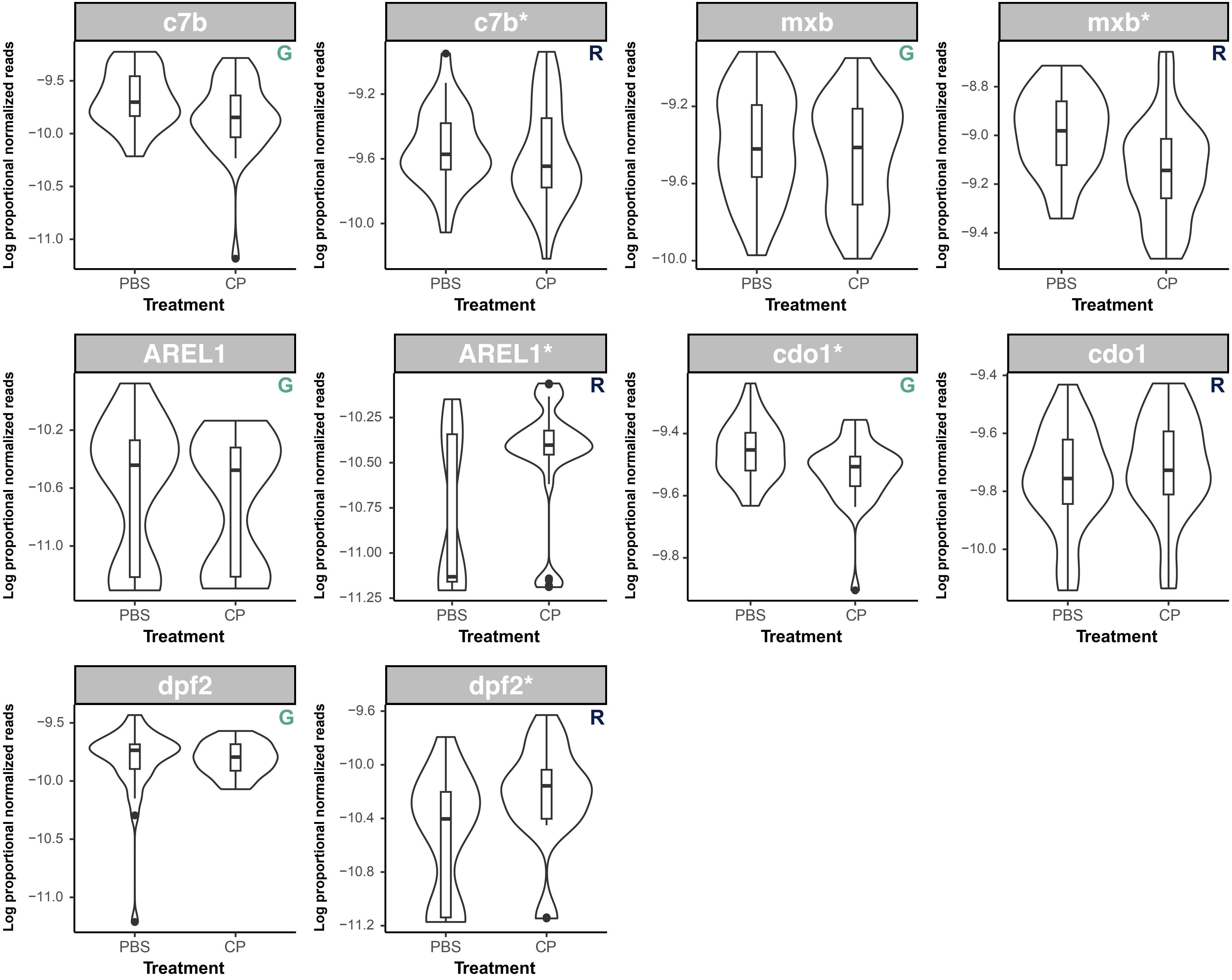
